# KLF4 promotes apoptosis evasion and PARP inhibitor resistance in BRCA2-mutated epithelial ovarian cancer

**DOI:** 10.64898/2026.08.23.746472

**Authors:** Esma Fera, Tiffany Zhang, Victoria M Grechukhina, Yong-Lian Zhu, Elena S Ratner, Z Ping Lin

**Affiliations:** Department of Obstetrics, Gynecology, and Reproductive Sciences, Yale University of School of Medicine, CT, USA

**Keywords:** KLF4, Epithelial ovarian cancer, Apoptosis, PARP inhibitor, Homologous recombination repair, DNA damage response

## Abstract

BRCA2-mutated epithelial ovarian cancer (EOC) is deficient in homologous recombination (HR) repair and hypersensitive to PARP inhibitors. However, BRCA2-mutated EOC frequently develops PARP inhibitor resistance and the underlying mechanisms involving apoptosis evasion remain poorly understood. In this study, our bioinformatic analysis of clinical transcriptomic datasets revealed that increased expression of KLF4, a zinc finger transcription factor, was strongly associated with high-grade serous EOC subtype and reduced overall survival of patients. Using isogenic EOC cells, we demonstrated that BRCA2 mutation led to pronounced KLF4 up-regulation by PARP inhibition in an ATM-dependent manner. Silencing of KLF4 and its target gene NR4A1 enhanced olaparib-induced apoptosis. Inhibition of anti-apoptotic effectors using the BH3-mimetic navitoclax, but not the SMAC-mimetic birinapant, selectively sensitized BRCA2-mutated EOC cells to olaparib. Furthermore, KLF4 silencing abrogated olaparib-induced BCL-w and BCL-xL, while olaparib-induced cIAP2 was attenuated only by NR4A1 silencing in BRCA2-mutated EOC cells. In vivo, combined treatment of navitoclax and olaparib synergized to impede the progression of BRCA2-mutated EOC xenografts and prolong mouse survival time. Collectively, our investigations discovered KLF4 as a regulatory hub of DNA damage response and apoptosis evasion in BRCA2-mutated EOC. These findings support targeting KLF4-driven anti-apoptotic pathways as a rational strategy to overcome PARP inhibitor resistance.

## Introduction

Ovarian cancer remains as one of the most lethal gynecologic malignancies among women. It was predicted that in 2026 around 21,010 women in the United States will be diagnosed with ovarian cancer and about 12,450 will die from the disease [1]. Epithelial ovarian cancer (EOC) is one of the histological subtypes accounting for the majority of ovarian malignancies. Approximately 10-15% of EOC patients are found to carry germ-line mutations in BRCA1 and BRCA2 genes [2]. BRCA1 and BRCA2 play a crucial role in the DNA repair mechanism via homologous recombination (HR) to restore genome stability following DNA damage. Therefore, mutations in BRCA1 and BRCA2 are major contributors to predisposition in EOC and importantly confer hypersensitivity to platinum-based chemotherapy and poly ADP-ribose polymerase (PARP) inhibitors [3]. Olaparib (Lynparza) is the first-in-class PARP inhibitor currently used as an important treatment strategy for patients diagnosed with BRCA-mutated or HR-deficient EOC [4].

Krüppel-like factor 4 (KLF4) is a member of the zinc finger-containing DNA-binding transcription factor subclass and regulates a variety of biological functions, including apoptosis, differentiation, proliferation, and reprogramming [5]. KLF4 consists of 483 amino acids with three C2H2 Krüppel-type zinc fingers in its C-terminal region. Previous sequence analysis showed that these zinc fingers share high similarity with other Krüppel-like family members [6,7]. Together with OCT4, SOX2, and c-MYC, KLF4 is one of the four Yamanaka transcription factors originally identified as sufficient to reprogram somatic cells into induced pluripotent stem cells [8], and has since been linked to tumorigenesis-related reprogramming [9]. However, KLF4 has been shown to operate as either an oncogene or a tumor suppressor depending on the context or cancer type [10]. For example, lung tumorigenesis was induced when KLF4 was downregulated due to USP10 loss [11]. KLF4 expression suppressed cell proliferation and promoted apoptosis in breast cancer [12,13]. Conversely, KLF4 was involved in cancer stem cell maintenance and enhanced migration and invasion of breast cancer [14]. Evidence also showed that KLF4 expression suppressed γ-irradiation-induced apoptosis in colorectal cancer cells [15]. Similarly, the role of KLF4 in ovarian cancer is also controversial. High KLF4 expression increased cisplatin resistance in ovarian cancer cells in vitro and is associated with platinum resistance and poor prognosis in patients [16]. On the other hand, it has been shown that the small molecule KLF4 inducer, APTO-253, sensitized ovarian cancer cell lines to paclitaxel and cisplatin [17]. Given the role of KLF4 in transcriptional regulation, a better understanding of its downstream signaling pathways in cell survival and apoptosis is necessary to settle these contradictory findings.

To date, a scarcity of studies has been conducted to investigate the involvement of KLF4 downstream effectors in the regulation of survival and apoptosis, especially in BRCA-mutated EOC. Thus, the objective of our study was to characterize the KLF4-mediated signaling pathway and examine its impact on the survival of BRCA2-mutated EOC cells in response to PARP inhibitor-induced DNA damage. Combining bioinformatic analyses of EOC patient datasets and experimental approaches with cell-based and in vivo models, we revealed the role of KLF4 in apoptosis evasion and PARP inhibitor resistance, as well as establishing a pragmatic strategy for enhancing the effectiveness of PARP inhibitor therapy for BRCA2-mutated EOC.

## Results

### BRCA2-mutated PEO1 cells exhibited increased KLF4 expression

We have previously demonstrated that BRCA2-mutated PEO1 cells expressed a higher level of KLF4 than BRCA2-wild type PEO4 cells [18]. Despite being isogenic from the same patients, PEO1 and PEO4 cells may have arisen from different ancestral cell populations due to their distinct growth characteristics and gene expression profiles. To investigate whether KLF4 expression was up-regulated as a result of BRCA2 mutation or homologous recombination (HR) deficiency, we established an isogenic PEO1 cell line PEO1-NR that was resistant to the PARP inhibitor niraparib. We sequenced the BRCA2 gene in PEO1-NR cells and confirmed that the nonsense mutation (TAG) was reverted to TTG that restored the reading frame of the BRCA2 gene (**Fig 1A**). Western blot analysis confirmed that PEO1-NR cells expressed a full length BRCA2 protein in contrast to a truncated BRCA2 protein expressed by PEO1 cells (**Fig 1B** and **Supplementary Fig 3**). In addition, PEO1-NR cells showed a reduced level of KLF4 compared to PEO1 cells (**Fig 1C**). To further substantiate this finding, we performed CRISPR mediated gene editing to abrogate the BRCA2 gene and HR repair in PEO1-NR cells. PEO1-NR-BRCA2-KO cells restored the heightened sensitivity to olaparib similar to PEO1 cells (**Fig 1D**). Furthermore, PEO1-NR-BRCA2-KO cells exhibited an increased KLF4 expression compared to PEO1-NR cells (**Fig 1E**). These results suggest that BRCA2 mutation or HR repair deficiency contributes to up-regulation of KLF4 in PEO1 cells.

**Figure 1.**
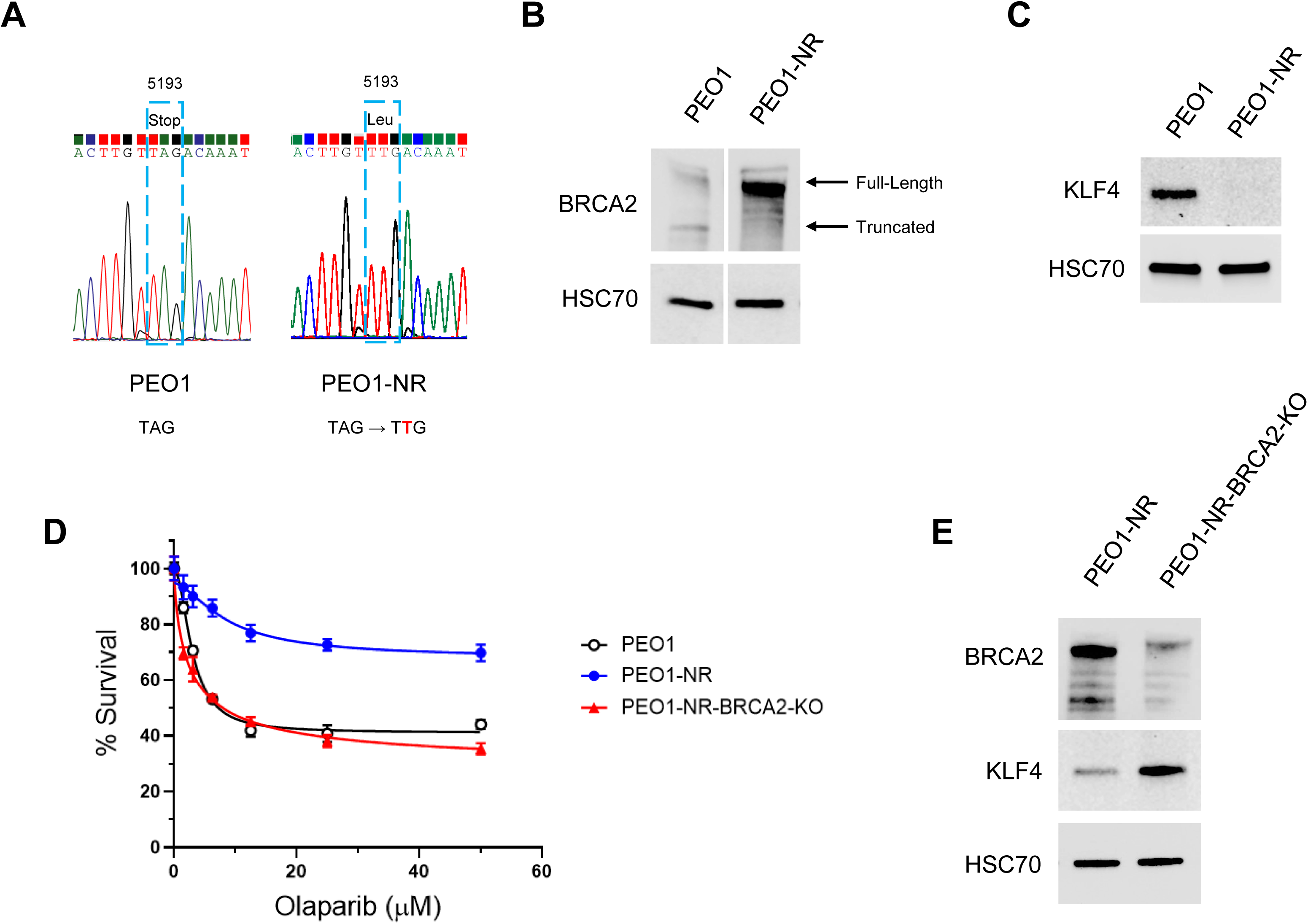
**A.** DNA sequencing for the mutation of the BRCA2 gene in PEO1 and PEO1-NR cells. PEO1 cells contain a nonsense mutation TAG (5193C>G) in the BRCA2 gene. PEO1-NR cells revert the mutation to TTG (5192A>T). The reverted mutation restored the reading frame of the BRCA2 gene. **B.** Western blot analysis of the BRCA2 protein expression in PEO1 and PEO1-NR cells. The truncated and full-length of BRCA2 proteins are shown. The HSC70 protein level was used as a loading control. **C.** Western blot analysis of the KLF4 protein expression level in PEO1 and PEO1-NR cells. The HSC70 protein level was used as a loading control. **D.** Cell viability/survival assay for the sensitivity of PEO1, PEO1-NR, and PEO1-NR-BRCA2-KO cells to a range of olaparib concentrations. Data are means ± SD (n=3). **E.** Western blot analysis of the BRCA2 and KLF4 protein expression levels in PEO1-NR and PEO1-NR-BRCA2-KO cells. The HSC70 protein level was used as a loading control.

### Loss of BRCA2 function increased the expression of KLF4 in response to DNA double strand breaks (DSBs) and IFN_γ_

Given that BRCA2 mutation or HR deficiency hinders the repair of DSBs, we investigated whether treatment with the PARP inhibitor olaparib induced KLF4 expression differentially in PEO1, PEO1-NR, and PEO1-NR-BRCA2-KO cells. Because KLF4 is an IFNγ- inducible gene [19], IFNγ was included in the treatments as a single agent and in combination with olaparib. Quantitative RT-PCR analysis showed that treatment with IFNγ had minimal effects on KLF4 mRNA levels but considerably enhanced olaparib-induced KLF4 mRNA. This effect was significantly greater in PEO1 and PEO1-NR-BRCA2-KO cells than in PEO1-NR cells (**Fig 2A**). To substantiate this finding, the mouse isogenic pair Brca2-WT and Brca2-KO ID8 cells were treated with increasing concentrations of olaparib. Quantitative RT-PCR analysis demonstrated that olaparib caused a significant and dose-dependent increase in Klf4 mRNA in Brca2-KO ID8 cells compared to Brca2-WT ID8 cells (**Fig 2B**). These results suggest that the expression of KLF4 is up-regulated by DSBs in the absence of BRCA2 function or HR repair in EOC cells.

**Figure 2.**
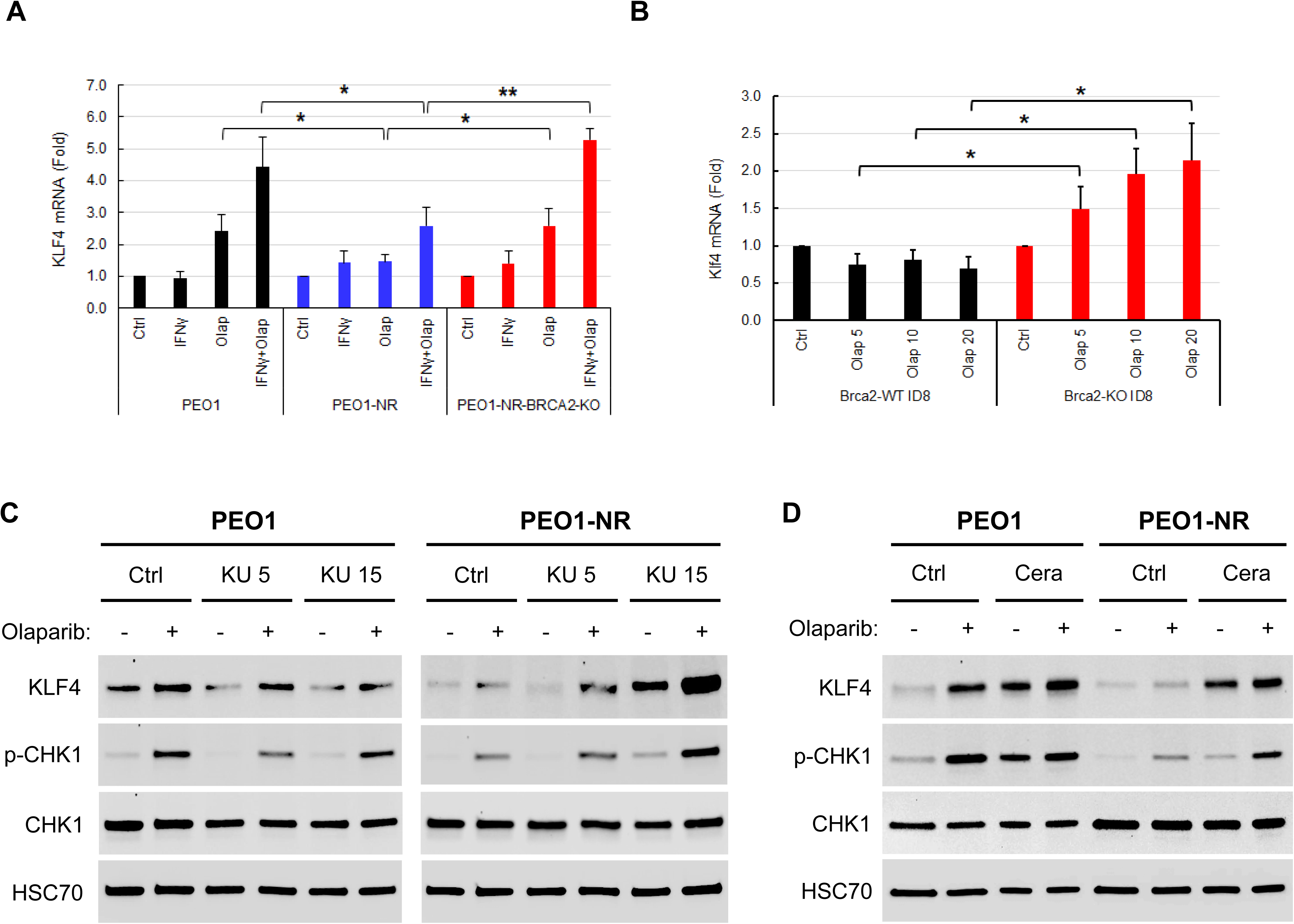
**A.** The effects of olaparib and IFNγ treatment on KLF4 mRNA expression in PEO1, PEO1- NR, and PEO1-NR-BRCA2-KO cells. Cells were treated with 10 µM olaparib, 10 ng/mL IFNγ, and both in combination for 24 hr. Total RNA was analyzed for KLF4 and GAPDH mRNA expression by quantitative RT-PCR. Fold increases in KLF4 mRNA relative to the vehicle-treated control in each cell line were shown. Data are ± SD (n=3). **B.** The effects of olaparib on Klf4 mRNA expression in mouse Brca2-WT and Brca2-KO ID8 cells. Cells were treated with a range of olaparib concentrations for 24 hr. Total RNA was analyzed for Klf4 and Gapdh mRNA expression by quantitative RT-PCR. Fold increases in Klf4 mRNA relative to the vehicle-treated control in each cell line were shown. Data are means ± SD (n=3). *, p<0.05. **, p<0.01. **C.** The effects of ATM inhibition on olaparib-induced KLF4 expression in PEO1 and PEO1-NR cells. Cells were treated with 5 or 15 μM of KU60019, 20 μM olaparib, or both in combination for 24 hr. Total protein was analyzed by western blotting for KLF4 and p-CHK1 expression levels. CHK1 and HSC70 protein levels were used as loading controls. **D.** The effects of ATR inhibition on KLF4 expression in PEO1 and PEO1-NR cells. Cells were treated with 1.25 μM of ceralasertib, 20 μM olaparib, or both in combination for 24 hr. Total protein was analyzed by western blotting for KLF4 and p-CHK1 expression levels. CHK1 and HSC70 protein levels were used as loading controls.

### ATM mediated KLF4 up-regulation by DSBs in BRCA2-mutated PEO1 cells

Given that ATM has been known to mediate DSB sensing and signaling [20], we sought to identify whether ATM plays a role in KLF4 up-regulation following PARP inhibitor-induced DSBs. PEO1 and PEO1-NR cells were treated with the ATM inhibitor KU60019 [21] in combination with olaparib. Western blot analysis demonstrated that in PEO1 cells, ATM inhibition caused a dose-dependent decrease in olaparib-induced levels of KLF4 (**Fig 2C** and **Supplementary Fig 4**). In PEO1-NR cells, however, ATM inhibition increased olaparib-induced levels of KLF4. Similarly to KLF4, ATM inhibition led to decreased levels of olaparib-induced CHK1 phosphorylation (p-CHK1) in PEO1 cells but elevated basal and olaparib-induced levels in PEO1-NR cells. Given that ATR and ATM play overlapping roles, PEO1 and PEO1-NR cells were treated with the ATR inhibitor ceralasertib [22] in combination with olaparib. Western blot analysis showed that ATR inhibition up-regulated KLF4 and p-CHK1 in PEO1 and, to a lesser extent, PEO1-NR cells (**Fig 2D** and **Supplementary Fig 4**). Collectively, these findings suggest that ATM predominantly mediates KLF4 up-regulation following DSBs in HR-deficient PEO1 cells. ATR serves as a backup mechanism to up-regulate KLF4 only when ATM function is absent in HR-proficient PEO1-NR cells.

### KLF4 up-regulation was correlated with high-grade serous subtype of EOC and decreased survival time in patients

To determine whether an increased KLF4 expression was associated with clinical features and prognosis of EOC in patients, we performed the Kaplan-Meier survival analysis to determine the KLF4 expression levels in relation to the survival of the EOC patients using the cBioPortal datasets. Six studies of serous ovarian cancer datasets consisting of 1047 patients were stratified based on KLF4 mRNA levels in tumor samples and compared for the overall survival of patients. The result showed that patients with high KLF4 expression exhibited a significantly shorter survival time than patients with lower KLF4 expression (logrank test, p< 0.01) (**Fig 3A**). In addition, the hazard ratio of the KLF4 high cohort was 1.241 compared to the KLF4 low cohort. Further analysis of clinical features of tumor samples showed that in the KLF4 high cohort, 73% of tumor samples displayed the high-grade serous subtype whereas only 20% of tumor samples showed the high-grade serous subtype in the KLF4 low cohort (**Fig 3B**). These differences were statistically significant (p<10^-10^). These findings suggest that KLF4 up-regulation is strongly associated with the high-grade subtype and reduced overall survival in patients.

**Figure 3.**
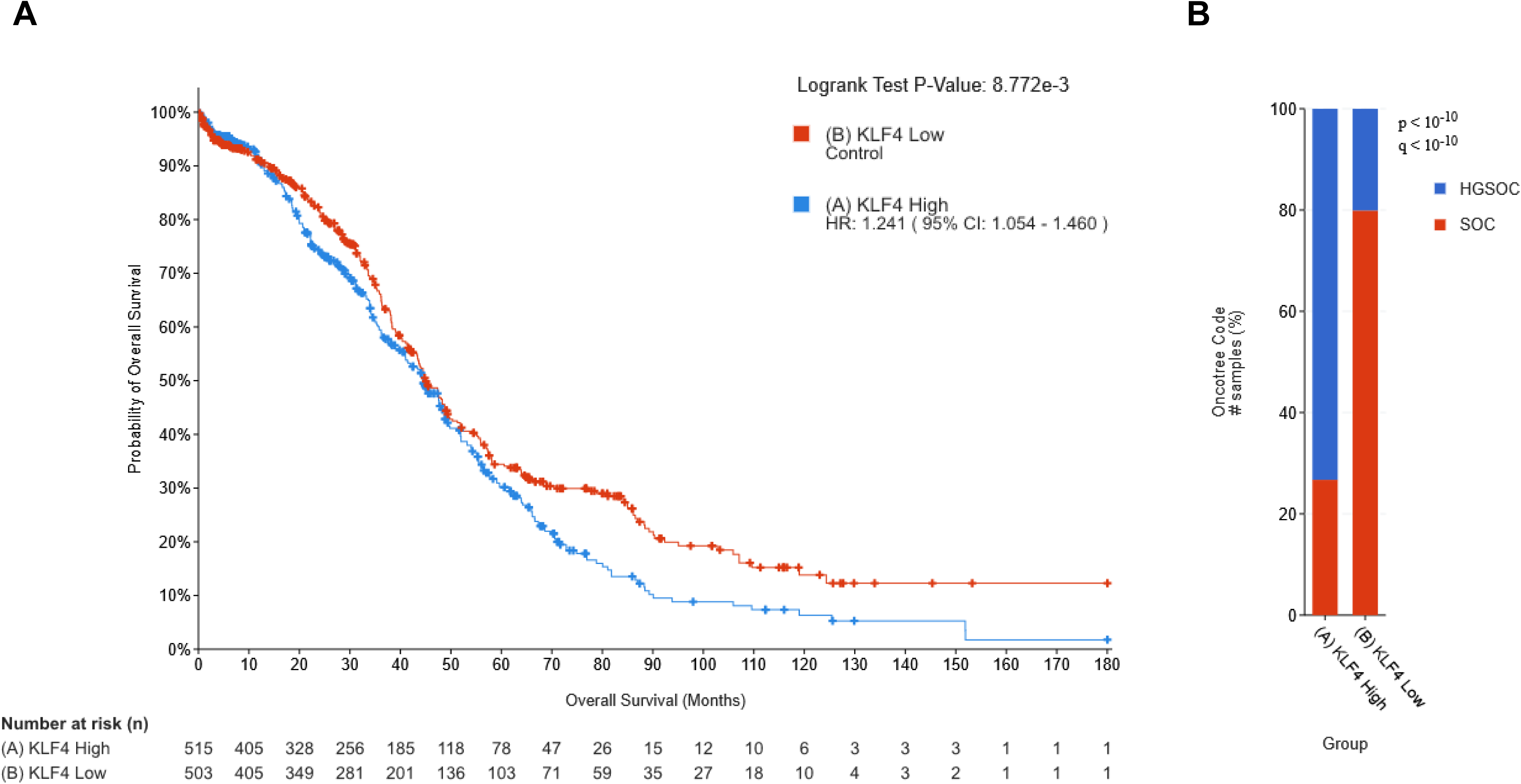
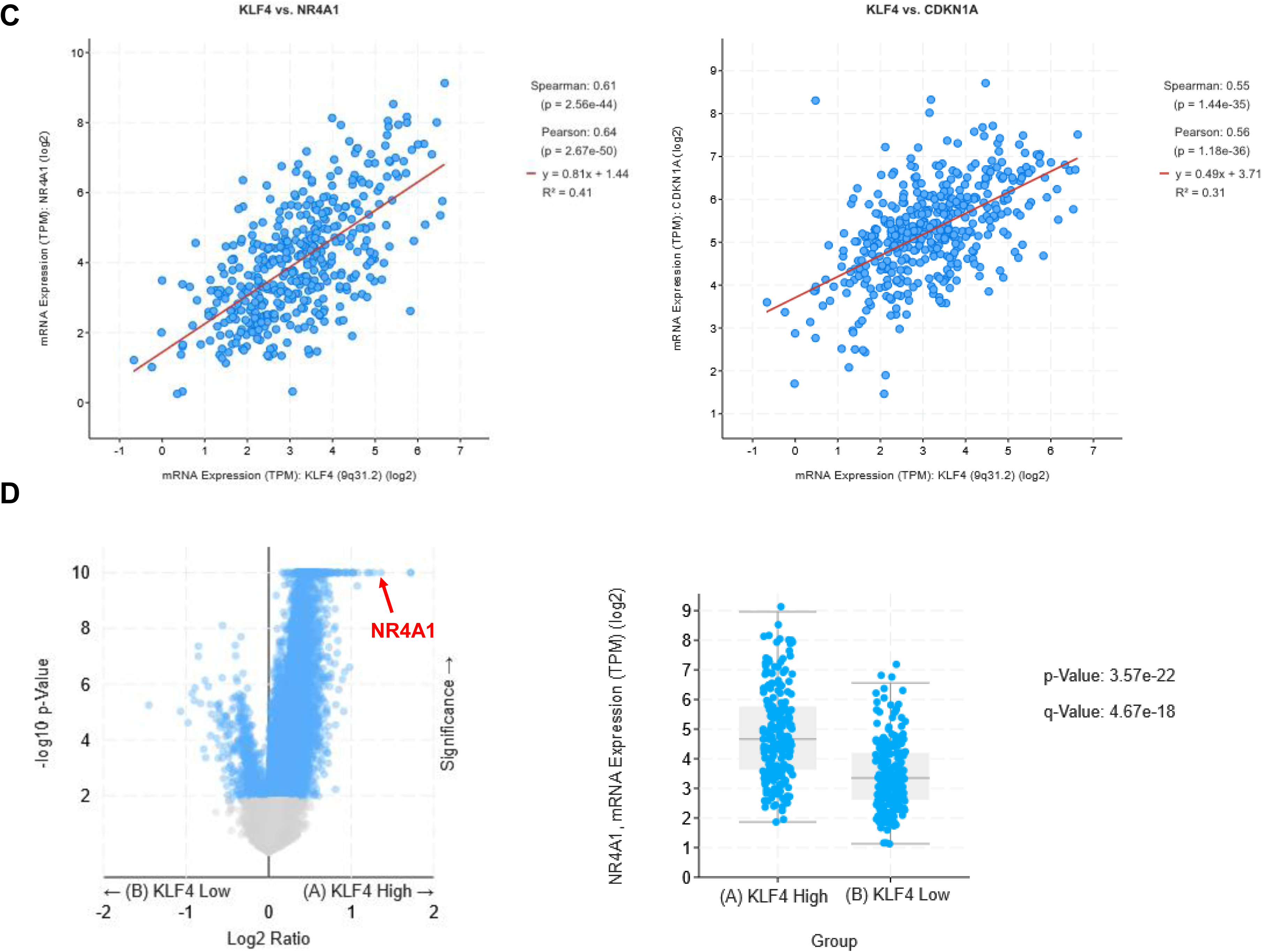
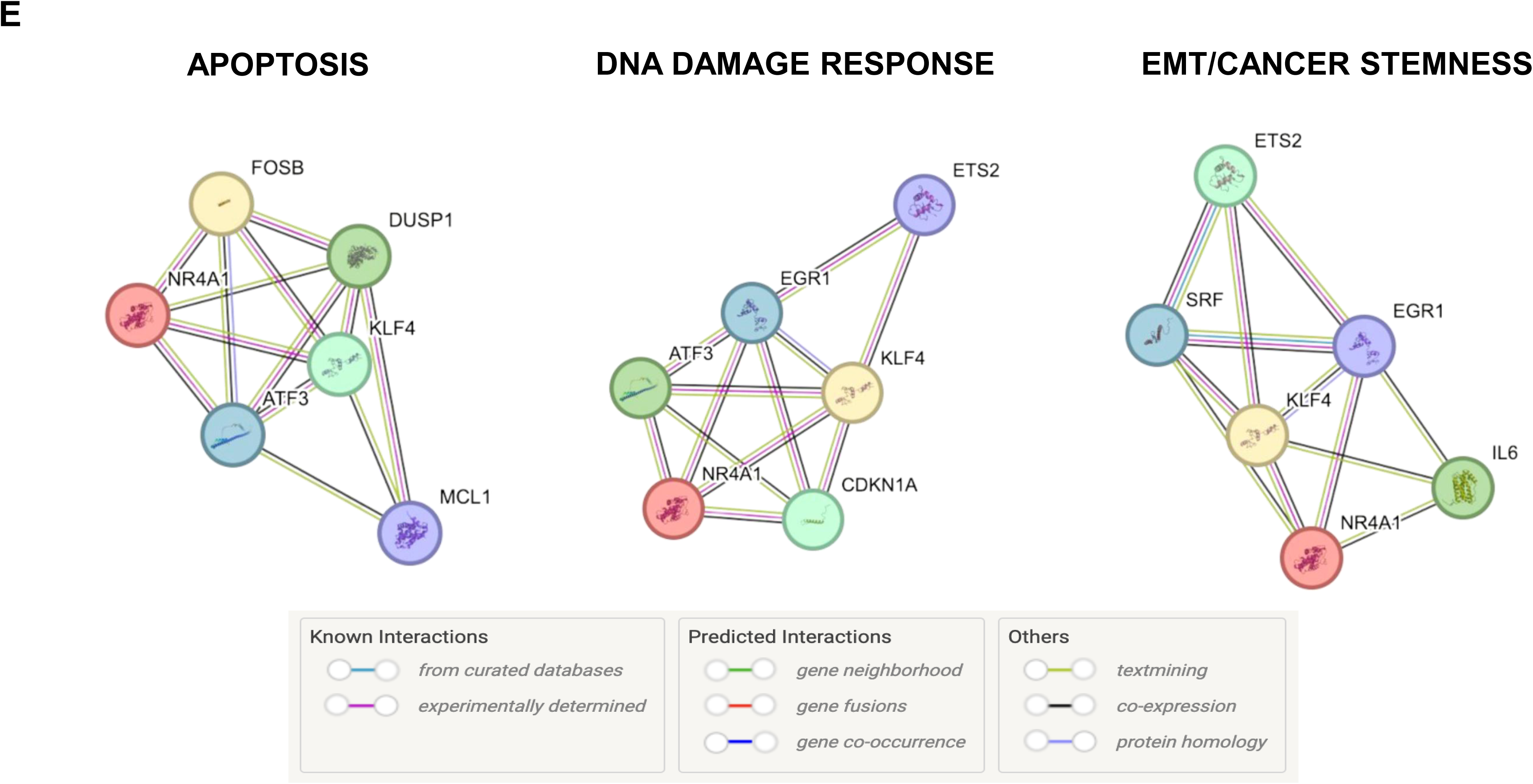
**A.** Kaplan-Meier survival analysis of ovarian cancer patients using cBioPortal datasets. Combined 1047 patients from 6 studies were stratified into KLF4 high and KLF4 low cohorts by the median KLF4 mRNA level. Statistical significance and hazard ratio (HR) are shown. **B.** KLF4 high and KLF4 low cohorts were compared for the prevalence of high-grade serous ovarian cancer (HGSOC) and serous ovarian cancer (SOC) subtypes in 1047 patients. Statistical significance (p-and q-values) is shown. **C.** Correlation analysis of KLF4 and NR4A1 expressions in high grade serous ovarian cancer patients (n=602) using the TCGA (GDC) dataset. The levels of KLF4 mRNA are plotted against the levels of NR4A1 or CDKN1A (p21) mRNA for each patient. A regression line is shown in red. Values of Spearman and Person correlations are also displayed. **D.** Volcano plot analysis of the association between KLF4 and NR4A1 mRNA expression (TPM) in high grade serous ovarian cancer. Patients were stratified into KLF4 high and KLF4 low cohorts by the median KLF4 mRNA level. NR4A1 is indicated by the arrow. The column scatter plot compared NR4A1 expression levels between KLF4 high and KLF4 low cohorts. Statistical significance (p-and q-values) is shown. **E.** STRING network analysis of KLF4 and NR4A1 grouped by functional modules in apoptosis, DNA damage response, and EMT/cancer stemness. KLF4, NR4A1, and associated proteins (nodes) are connected with predicted functional associations or physical interactions (edges). Types of interactions are indicated in colors in the legend.

### Bioinformatic analyses of top KLF4-correlated genes and protein-protein interaction networks

To investigate how KLF4 expression correlated with transcriptional programs driving ovarian cancer phenotypes, we analyzed mRNA co-expression patterns using the cBioPortal dataset of high-grade serous ovarian cancer from 602 patients (TCGA, GDC). In the scatter plot, KLF4 showed strong positive correlations with many genes, with NR4A1 being the top association (Spearman: 0.61, p=2.57 × 10^-44^) (**Fig 3C** and **Supplementary Table 1**). Additional correlated genes included CDKN1A (p21) (Spearman: 0.55, p=1.44 × 10^-35^) and MCL1 (Spearman: 0.48, p=2.12 × 10^-26^). In addition, the volcano plot and stratified group analyses showed that NR4A1 was expressed in positive and significant association with KLF4 expression (**Fig 3D**).

KLF4 has been implicated in regulating stemness/mesenchymal properties, DNA damage responses, and apoptotic signaling [10,23,24]. To characterize potential relationships among these genes, protein– protein interaction analysis was performed using the STRING database. STRING integrates known and predicted protein associations derived from multiple sources including experimental interaction datasets, curated pathway databases, gene co-expression, genomic context predictions, and literature text mining [25]. The resulting network visualization revealed that KLF4-associated genes can be clustered into three major functional modules: apoptosis (NR4A1, DUSP1, ATF3, MCL1, and FOSB), DNA damage response (NR4A1, CDKN1A, EGR1, ETS2, and ATF3), and epithelial to mesenchymal transition (EMT)/cancer stemness (NR4A1, IL6, SRF, EGR1, and ETS2) (**Fig 3E**). Noticeably, many of these genes were shared among these functional modules. These networks suggest that KLF4 may contribute to the regulation of three interlinked processes in cancer.

### KLF4 silencing reduced target gene expression and enhanced apoptosis in BRCA2-mutated PEO1 cells

Given the strong correlation of KLF4 with NR4A1 and CDKN1A (p21) observed in bioinformatic analyses, we examined whether KLF4 contributed to the induction of these genes following PARP inhibition by olaparib. In PEO1 cells, olaparib caused a considerable increase in the expression of KLF4, NR4A1, and p21 mRNA. Silencing of KLF4 significantly reduced olaparib-induced KLF4, NR4A1, and p21 mRNA (**Fig 4A**). Due to wild type BRCA2 and functional HR status, PEO1-NR cells showed minimal induction of these genes by olaparib. KLF4 silencing only significantly reduced olaparib-induced p21 mRNA. Because KLF4-associated genes were involved in apoptosis and DNA damage response, we evaluated whether silencing of KLF4, NR4A1, or p21 expression influenced apoptosis in response to PARP inhibition. In PEO1 cells, silencing of KLF4 or NR4A1 led to a significant and time-dependent increase in olaparib-induced apoptosis. In PEO1-NR cells, KLF4 silencing had no effect on olaparib-induced apoptosis but NR4A1 silencing significantly increased apoptosis only at 6 hr. In contrast, p21 silencing did not increase olaparib-induced apoptosis across all time points in both PEO1 and PEO1-NR cells (**Fig 4B**). Collectively, these findings suggest that PARP inhibition up-regulates KLF4 and its target genes NR4A1 and p21 in the absence of BRCA2 or HR function. Both KLF4 and NR4A1 function to suppress PARP inhibition-induced apoptosis in PEO1 cells, whereas only NR4A1 serves to suppress apoptosis in PEO1-NR cells.

**Figure 4.**
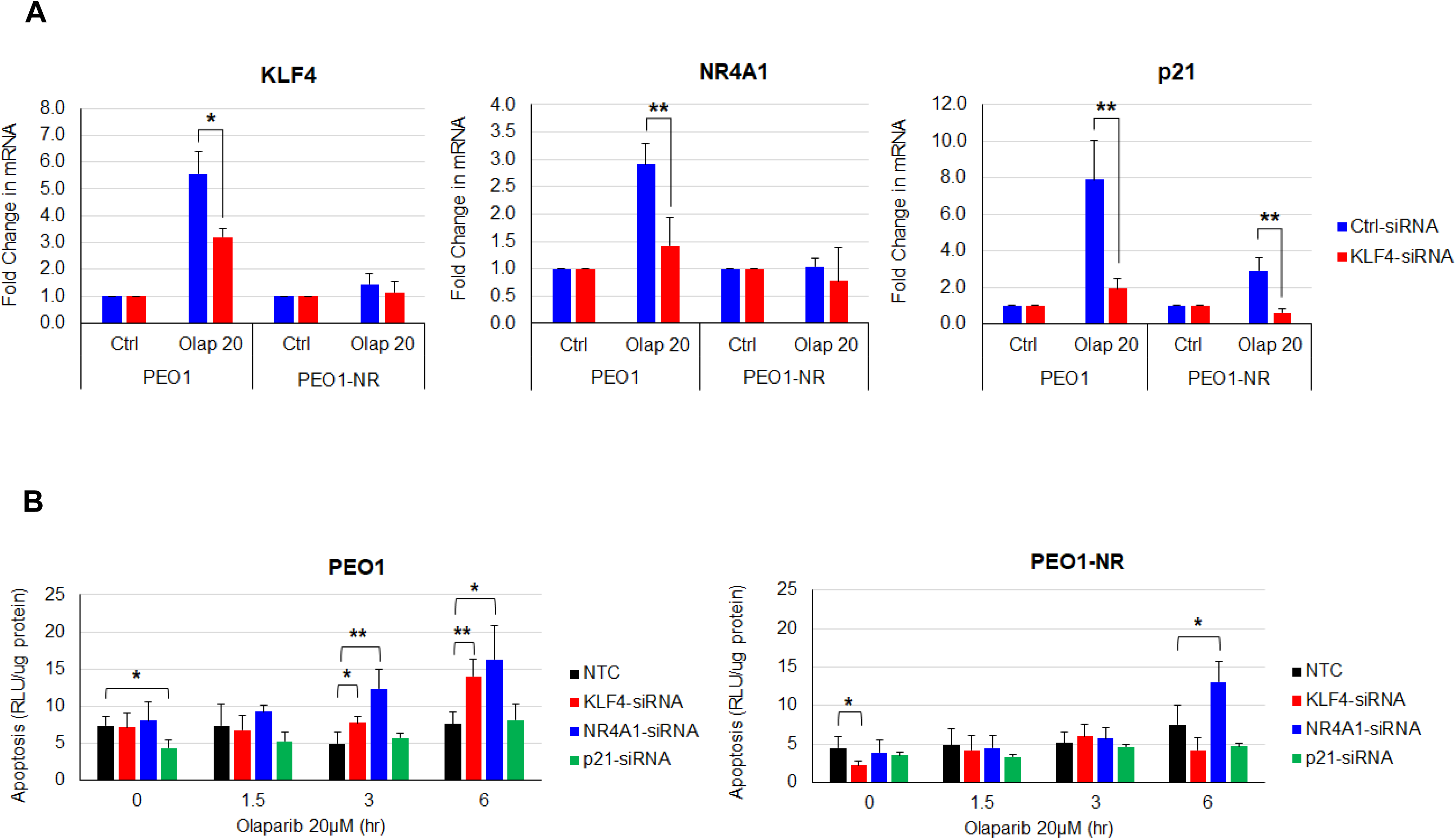
**A.** The effects of KLF4 silencing on KLF4-target gene expression in PEO1 and PEO1-NR cells. Cells were transfected with KLF4 siRNA for 24 hr and then treated with 20 µM olaparib for additional 24 hr. Total RNA was analyzed by quantitative RT-PCR to determine the fold change in KLF4, NR4A1, and p21 mRNA levels compared to vehicle-treated control for each cell line. **B.** The effects of KLF4, NR4A1, and p21 silencing on apoptosis in PEO1 and PEO1-NR cells. Cells were transfected with KLF4-, NR4A1-, or p21-siRNA for 42 hr. Thereafter, cells were treated with 20 µM olaparib for 1.5, 3, or 6 hr. Cell lysates were analyzed by Caspase 3/7 assay to determine the level of apoptosis expressed as RLU/µg protein. Data are means ± SD (n=3). *, p<0.05. **, p<0.01.

### BCL2 inhibitors enhanced olaparib-induced apoptosis in BRCA2-mutated PEO1 cells

Because anti-apoptotic BCL2 proteins play an important role in suppressing apoptosis [26], we explored whether pharmacological blockade of these BCL2 proteins selectively enhanced PARP inhibition-induced apoptosis in BRCA2-mutated PEO1 cells over BRCA2-wild type PEO1-NR cells. We performed the cell viability/survival assay to evaluate the efficacy of combining olaparib and a BCL2 inhibitor in PEO1 and PEO1-NR cells. BH3 mimetic drugs [27] navitoclax [28], sabutoclax [29], venetoclax [30], and A1210477 [31] that target a single or multiple BCL2 protein family members, BCL2, BCL-xL, BCL-w, and MCL1 (**Supplementary Table 2**) were used. In PEO1 cells, the combination of olaparib and navitoclax strongly reduced cell survival showing high Loewe and ZIP synergy scores, while the combination exhibited low synergy scores in PEO1-NR cells (**Fig 5A** and **5B**). In contrast, the combination of olaparib with sabutoclax, venetoclax, or A1210477 showed a comparable efficacy in reducing the survival of PEO1 and PEO1-NR cells (**Supplementary Fig 1A** and **Supplementary Fig 2A**), as judged by low synergy scores (**Supplementary Fig 1B** and **Supplementary Fig 2B**). To corroborate the synergistic efficacy of the olaparib-navitoclax combination, the apoptosis assay was carried out. In PEO1 cells, navitoclax alone at 3 and 6 µM produced considerable levels of apoptosis but significantly enhanced olaparib-induced apoptosis. In contrast, navitoclax at these concentrations did not significantly enhance apoptosis when combining with olaparib in PEO1-NR cells (**Fig 5C**).

**Figure 5.**
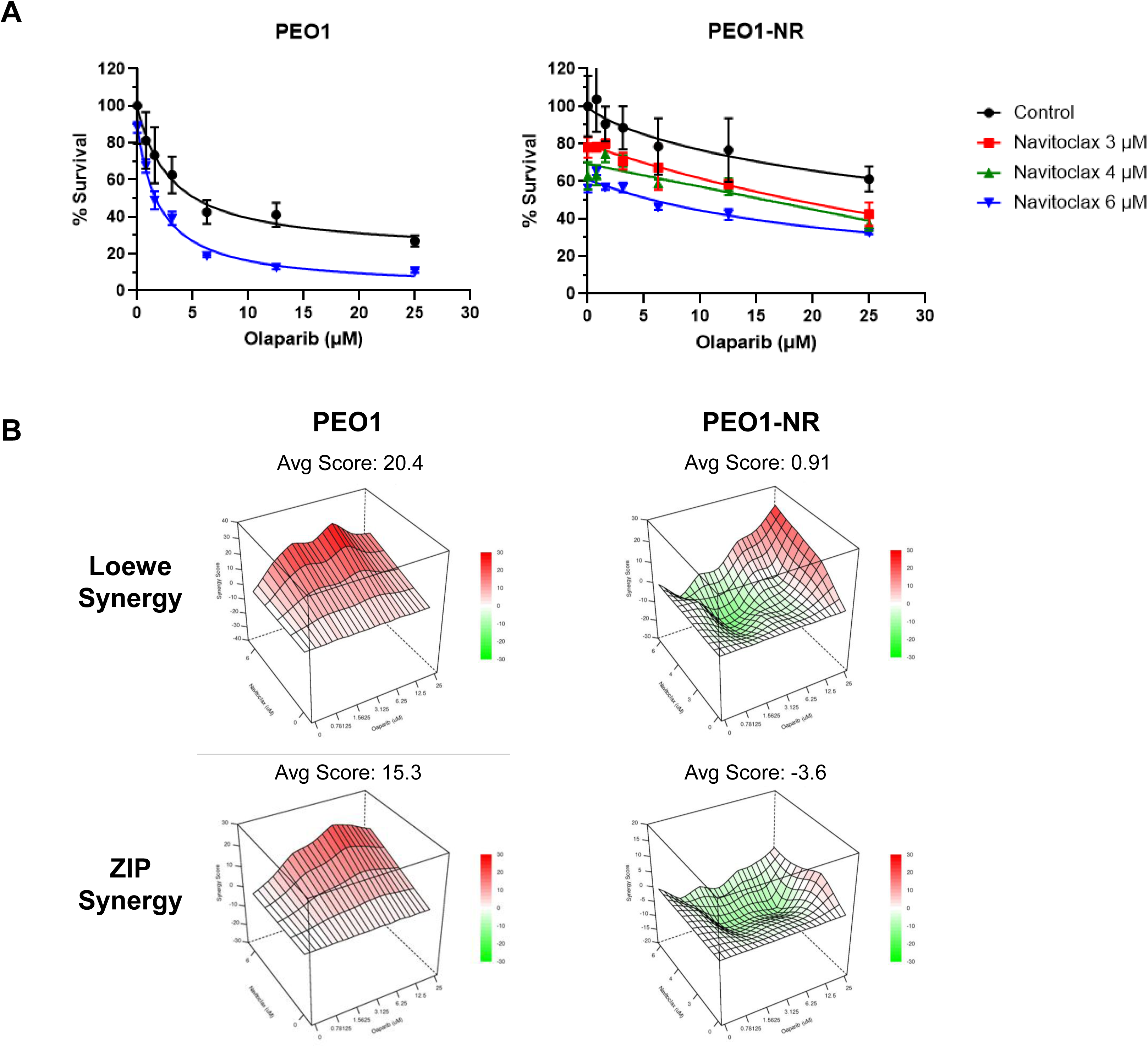
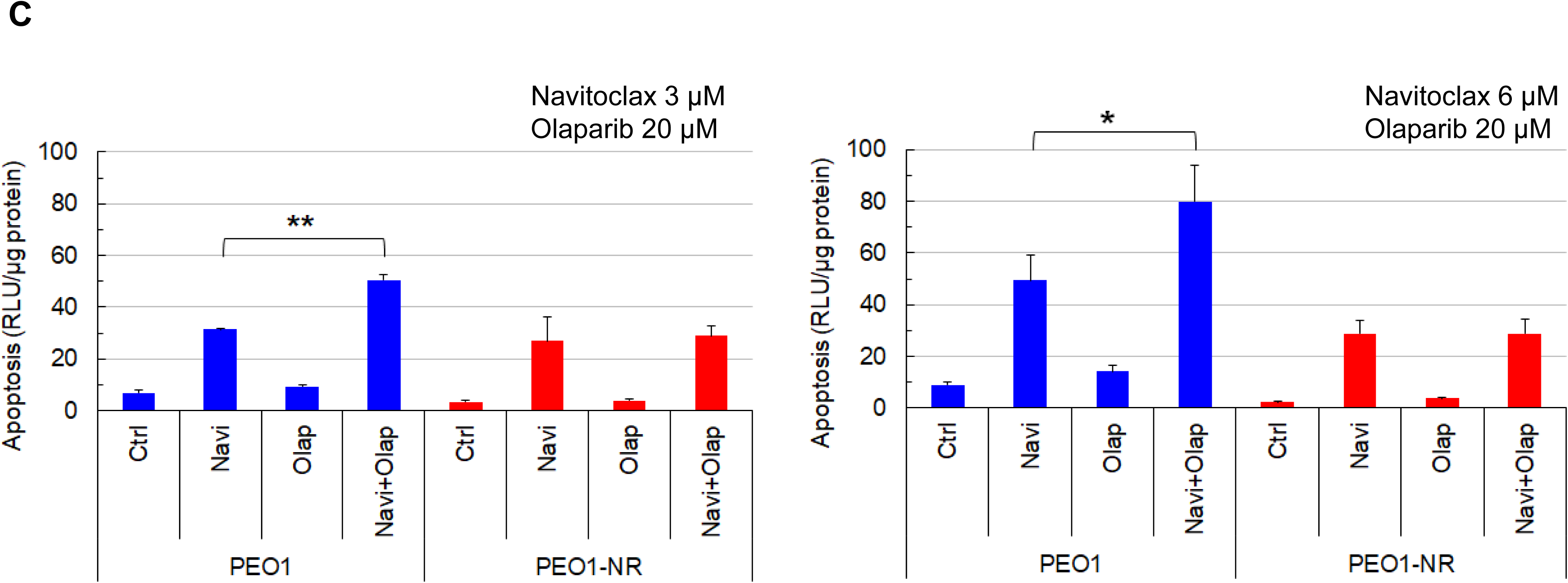
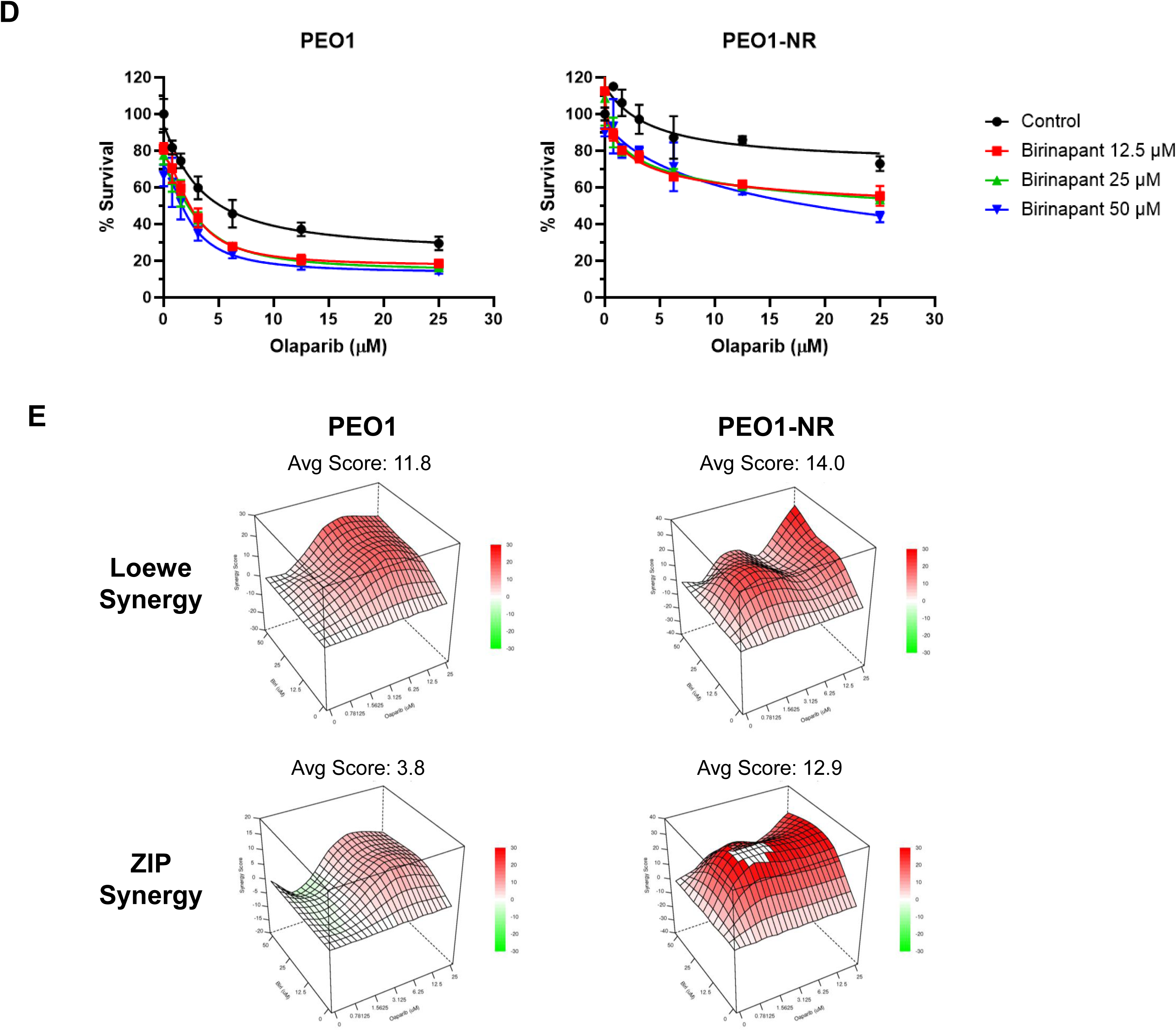
The effects of navitoclax on the sensitivity of PEO1 and PEO1-NR cells to olaparib. **A.** PEO1 cells were treated with 3 µM navitoclax and PEO1-NR cells were treated with 3, 4, or 5 µM navitoclax in combination with a range of concentrations of olaparib for 72 hr. Cell viability was determined by the MTS cytotoxicity assay and expressed as % survival. Data are means ± SD (n=3-4). **B.** Data of cell viability were used to determine the synergy score using the SynergyFinderPlus website. Synergy scores using Loewe and ZIP methods are shown to evaluate the synergistic effect of the combination of olaparib and navitoclax. Larger scores indicate greater synergistic effects between two drugs. **C.** Cells were treated with 3 or 6 µM navitoclax in combination with 20 µM olaparib for 24 hr. Cell lysates were analyzed by Caspase 3/7 assay to determine the level of apoptosis expressed as RLU/µg protein. Data are means ± SD (n=3). *, p<0.05. **, p<0.01. **D.** PEO1 and PEO1-NR cells were treated with 12.5, 25, and 50 µM birinapant in combination with a range of concentrations of olaparib for 72 hr. Cell viability was determined by the MTS cytotoxicity assay and expressed as % survival. Data are means ± SD (n=3-4). **E.** Synergy scores using Loewe and ZIP methods are shown to evaluate the synergistic effect of the combination of olaparib and birinapant.

Furthermore, we investigated whether pharmacological blockade of anti-apoptotic cIAP proteins had differential effects on olaparib-induced apoptosis between PEO1 and PEO1-NR cells. Birinapant is a Smac mimetic drug that inhibits cIAP1 and cIAP2 [32]. In both PEO1 and PEO1-NR cells, the combination of olaparib and birinapant led to similarly reduced cell survival, as demonstrated by moderate Loewe and ZIP synergy scores (**Fig 5D** and **E**). Taken together, these results suggest that BRCA2-mutated PEO1 cells up-regulate anti-apoptotic BCL2 proteins, thereby mitigating apoptosis in response to PARP inhibition.

### Characterization of KLF4-regulated anti-apoptotic effector proteins

To further characterize the KLF4-NR4A1 axis, we used the Gene Regulatory Network Database (GRAND) to identify the regulatory relationships in PEO1 cells. Within our apoptotic gene network, GRAND analysis revealed several regulator-target connections, the core of which is the KLF4 to NR4A1 connection, with an edge weight of 2.69, indicating qualitative support for this regulator-target connection in this model (**Fig 6A**). Additional regulatory edges identified include NR4A1 to cIAP2 (edge weight 2.91), KLF4 to BCL-w (2.33), KLF4 to BCL-xL (1.76), and KLF4 to MCL1 (2.08). Together, these bioinformatic results inform the association and directionality of our proposed apoptotic pathway involving KLF4 and NR4A1. To experimentally evaluate this pathway in our model, we examined whether olaparib-induced KLF4 protein expression was accompanied by corresponding changes in its predicted downstream targets. In PEO1 cells, increased concentrations of olaparib markedly induced expression of γ-H2AX (**Fig 6B** and **Supplementary Fig 5**), a marker of DSBs. In contrast, olaparib treatment in PEO1-NR cells induced a mild increase in γ-H2AX expression, suggesting that wild type BRCA2 or functional HR repair reduces olaparib-induced DSBs. PEO1 cells also demonstrated a pronounced dose-dependent increase in KLF4 expression, whereas PEO1-NR cells showed no increase in KLF4 expression with increasing olaparib concentrations. Of the BCL-2 family proteins, BCL-xL and BCL-w were most notably induced by increased olaparib concentrations in PEO1 cells, with no visible induction present in PEO1-NR cells. In contrast, no apparent trends in BCL2 or MCL1 expression were observed in either PEO1 or PEO1-NR cells. Notably, NR4A1 expression was higher at baseline in PEO1-NR cells than in PEO1 cells and largely unchanged across the olaparib dose range (**Fig 6B**). Among cIAP family proteins, cIAP1 showed no consistent trend across PEO1 and PEO1-NR cells. In contrast, cIAP2 exhibited dose-dependent induction in PEO1 cells but a high baseline expression in PEO1-NR cells. Given that olaparib-induced NR4A1 was not observed at the 24 hr time point and NR4A1 is known as an immediate early-response gene [33], we examined its temporal response to olaparib in PEO1 cells at the early phase of olaparib treatment. The time-course study showed that olaparib-induced NR4A1 peaked at 1.5 hr followed by a decline as early as 3 hr. In addition, this early induction of NR4A1 was completely abrogated by KLF4 silencing (**Fig 6C**). Because of the observed association between olaparib-induced KLF4 expression and the induction of anti-apoptotic proteins, we next investigated the impacts of KLF4 silencing on BCL2 and cIAP proteins. KLF4-silenced PEO1 cells showed a prominent reduction in olaparib-induced BCL-w and BCL-xL protein expression. Bioinformatic analysis of high-grade serous EOC also showed that besides MCL1, BCL-w expression was the most significantly elevated in the KLF4 high cohort among BCL2 family members (**Supplementary Table 3**). In PEO1-NR cells, KLF4 silencing appeared to enhance the olaparib-induced BCL-w and overall BCL-xL response, suggestive of unknown regulatory mechanisms compensating for KLF4 loss (**Fig 6D**). No apparent changes in cIAP2 expression were observed following KLF4 silencing in PEO1 and PEO1-NR cells (**Fig 6E**). However, olaparib-induced cIAP2 expression was abrogated by NR4A1 silencing in PEO1 cells (**Fig 6F**). Together, these findings suggest that olaparib-induced BCL-w and BCL-xL expression is KLF4-dependent in PEO1 cells. Despite KLF4 directly regulates NR4A1 induction, NR4A1 can mediate olaparib-induced cIAP2 expression in a manner bypassing KLF4.

**Figure 6.**
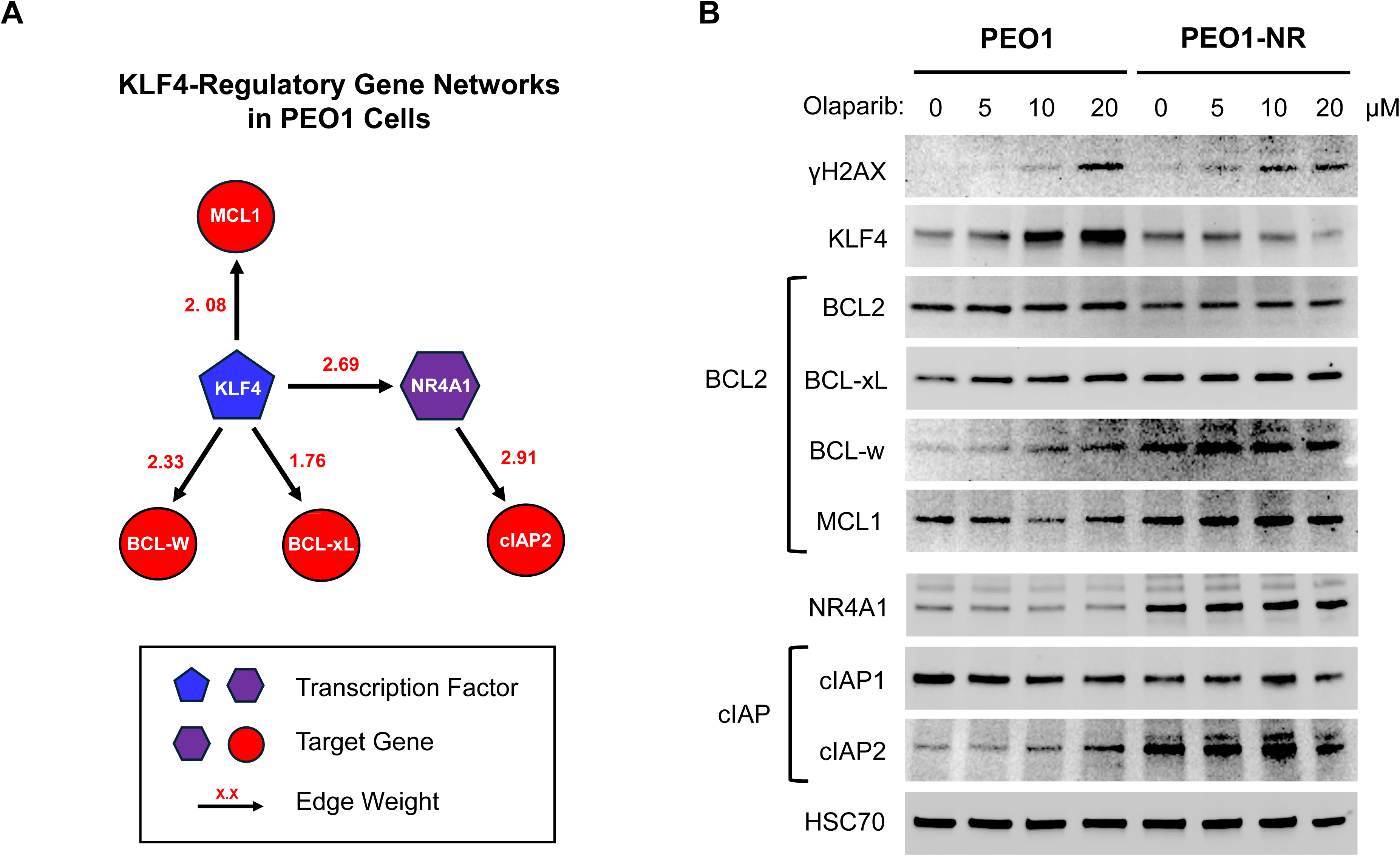
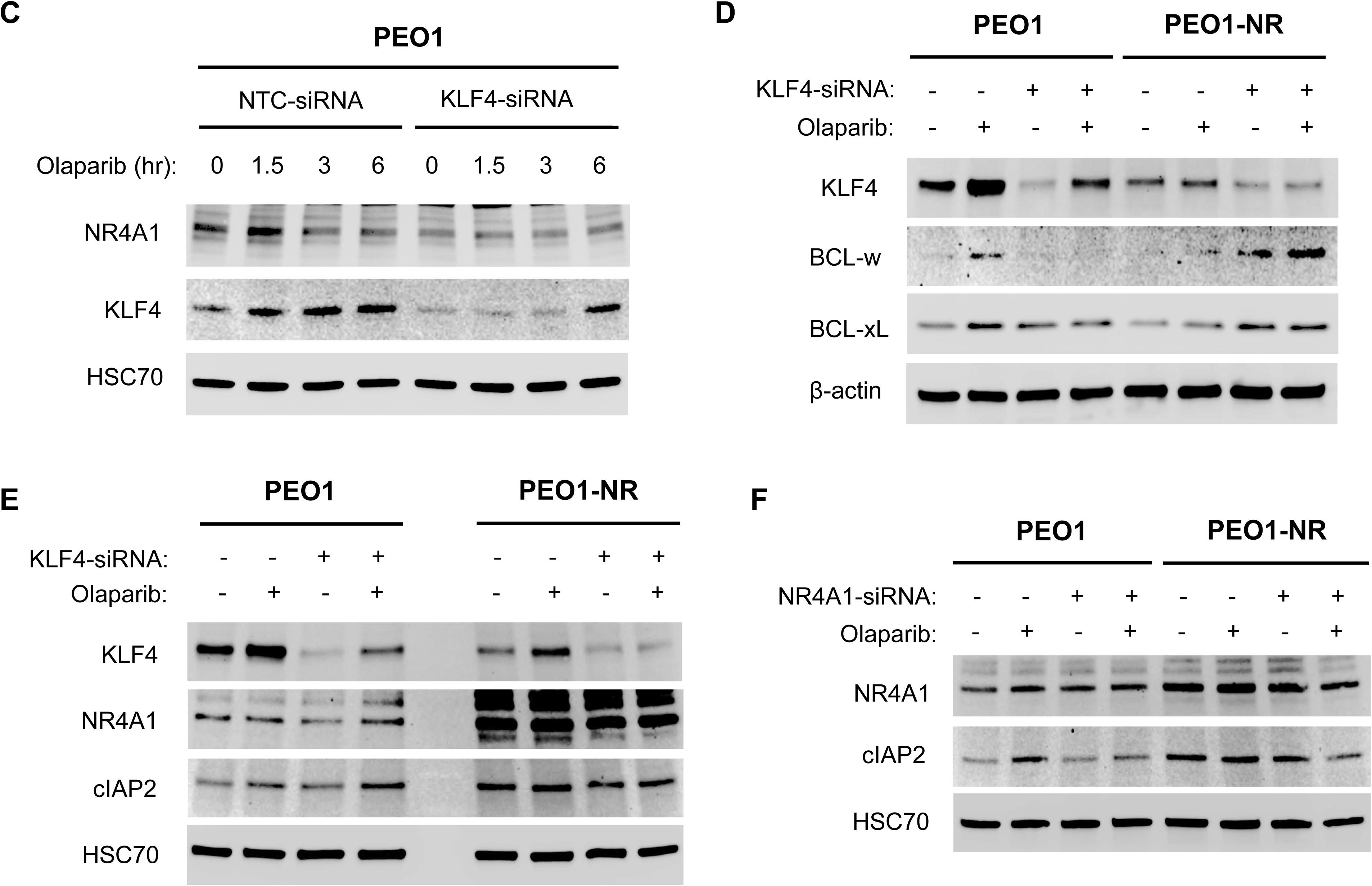
The effects of olaparib and silencing of KLF4 or NR4A1 on protein levels in the KLF4-and NR4A1-asssociated pathways in PEO1 and PEO1-NR cells. **A**. Bioinformatic analysis using the GRAND database depicts network connectivity among KLF4, NR4A1, BCL-w, BCL-xL, MCL1, and cIAP2 proteins via edge weights in PEO1 cells. **B**. PEO1 and PEO1-NR cells were treated with 0, 5, 10, or 20 μM olaparib for 24 hr. Total protein was analyzed by western blotting for γH2AX, KLF4, NR4A1, BCL2 family, and cIAP family protein levels. The HSC70 protein level was used as a loading control. **C.** PEO1 cells were transfected with non-targeted siRNA control or KLF4-siRNA. Cells were treated with 20 μM olaparib for 1.5, 3, or 6 hrs. Total protein was analyzed by western blotting for NR4A1 and KLF4 protein levels. The anti-NR4A1 antibody (RRID: AB_3750353) was used in this blot. The HSC70 protein level was used as a loading control. **D**. PEO1 and PEO1-NR cells were transfected with non-targeted siRNA control or KLF4-siRNA and treated with 20 μM olaparib for 24 hr. Total protein was analyzed by western blotting for KLF4, BCL-w, BCL-xL protein levels. The anti-BCL-w antibody (RRID:AB_10983656) was used in this blot. The β-actin protein level was used as a loading control. **E.** PEO1 and PEO1-NR cells were transfected with non-targeted siRNA control or KLF4-siRNA and treated with 20 μM olaparib for 24 hr. Total protein was analyzed by western blotting for KLF4, NR4A1, and cIAP2 protein levels. The HSC70 protein level was used as a loading control. **F.** PEO1 and PEO1-NR cells were transfected with non-targeted siRNA control or NR4A1-siRNA and treated with 20 μM olaparib for 24 hr. Total protein was analyzed by western blotting for NR4A1 and cIAP2 protein levels. The HSC70 protein level was used as a loading control.

### Navitoclax enhanced the effectiveness of olaparib to delay the progression of PEO1 xenografts in mice

Because navitoclax and olaparib synergistically reduced the survival of PEO1 cells in cell-based assays, we sought to evaluate the efficacy of this combination to treat PEO1 tumor xenografts in vivo. SCID-Beige mice were implanted with PEO1ip cells [34] and treated with vehicle, navitoclax, olaparib, and both drugs in combination for 4 weeks. The results showed that the control group of mice reached the maximal body weight around day 50 followed by a steady decline (**Fig 7A**). The body weight fell below the original body weight on day 80. The navitoclax group also reached the maximal weight, albeit lower than the control group, around day 50. The body weight fell below the original body weight on day 94. The olaparib group reached the maximal weight spanning around day 50 and maintained at a similar level up to day 120. In contrast, the combination group exhibited a continuous increase in body weight beyond day 120. The observation implies that both olaparib and olaparib plus navitoclax groups delay the onset and progression of PEO1ip xenografts in mice. Furthermore, Kaplan-Meier survival analysis demonstrated that control, navitoclax, and olaparib group reached the median survival times of 94, 94, and 140 days, respectively (**Fig 7B**). The olaparib plus navitoclax group had a median survival time of more than 175 days because 50% of mice continued to survive beyond 200 days. Navitoclax group was not significantly different from the control group. In contrast, olaparib (p<0.05) and olaparib plus navitoclax (p<0.01) groups showed a significantly longer time than the control group. Importantly, the survival time of the olaparib plus navitoclax group was significantly longer than the olaparib group (p<0.01). The hazard ratios of navitoclax, olaparib, and olaparib plus navitoclax groups to the control group were 0.696, 0.294, and 0.173, respectively (**Fig 7C**). Taken together, these findings suggest that treatment with navitoclax has no effects but significantly augments the effectiveness of olaparib to deter the progression of BRCA2-mutated EOC xenografts and extend the survival time of mice.

**Figure 7.**
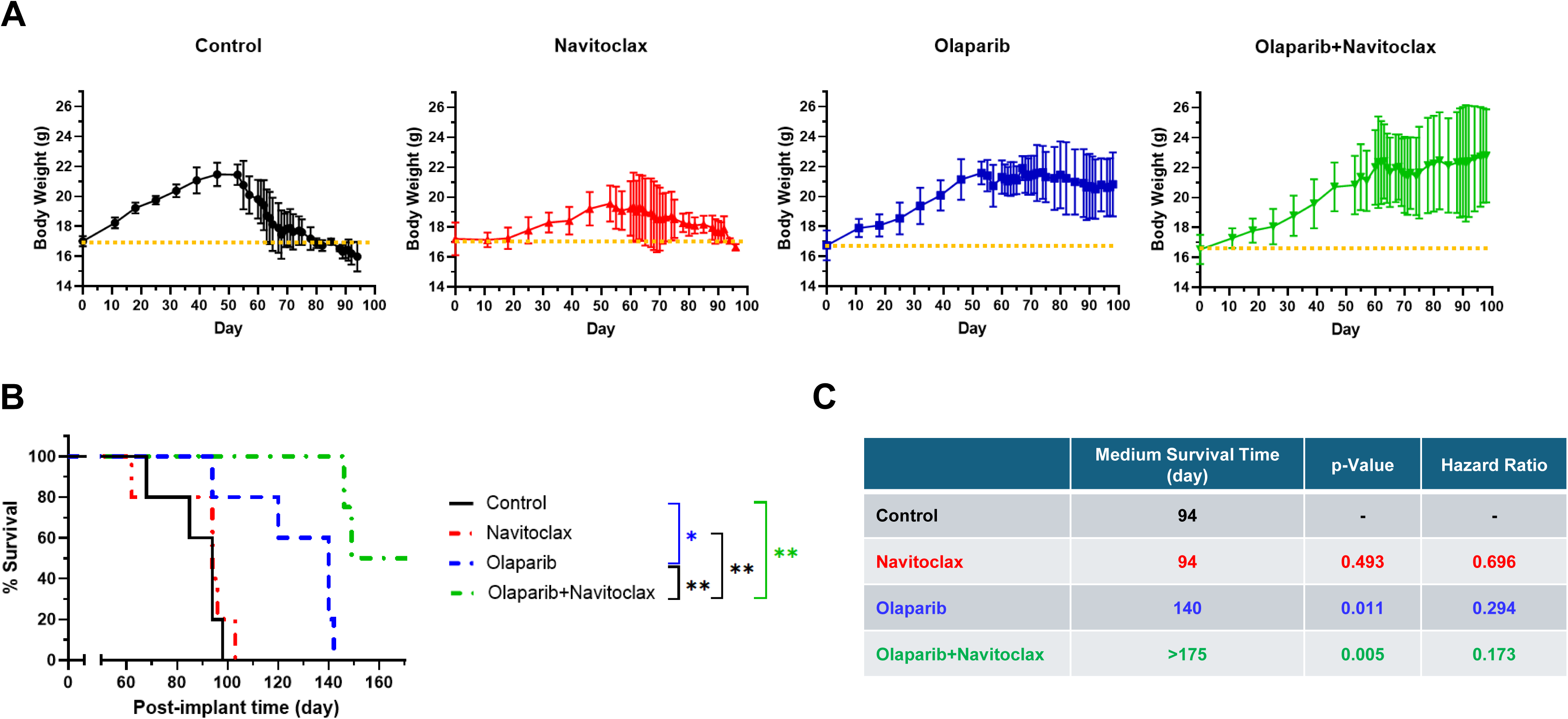
The effects of the combination of navitoclax and olaparib on the survival of SCID-Beige mice bearing peritoneal PEO1ip xenografts. Mice (n=5, except for the olaparib+navitoclax group, n=4) were inoculated ip with PEO1ip cells and treated ip with vehicle, olaparib, navitoclax, or olaparib plus navitoclax daily (except for weekends) for 4 weeks. **A.** The body weight of mice was measured to monitor cachexia caused by the progression of PEO1ip xenografts. The body weight of mice for the first 100 days following implantation was shown. Data are means ± SD (n=4-5). **B.** The Kaplan-Meier survival curves of mice were determined using BCS2 as the survival endpoint. *, p<0.05. **, p<0.01. **C.** The median survival times for each treatment group, p values compared to the control group, and the hazard ratios (log-rank) of each treatment group to the control group are shown.

## Discussion

PARP inhibitors such as olaparib have significantly improved outcomes for patients with BRCA-mutated EOC, however both intrinsic and acquired resistance still pose major clinical challenges [35]. Pathways that influence PARP inhibitor sensitivity or resistance are numerous and can arise through multiple adaptive mechanisms. Many studies have shown how certain mechanisms influence PARP inhibitor sensitivity including checkpoint rewiring (ATR–CHK1) [36], up-regulation of the NOTCH signaling [37], BRCA1/2 reversion mutations that restore HR [38], drug-efflux–mediated resistance via ABCB1/P-glycoprotein induction [39], and more recently, metabolic rewiring through activation of de novo lipid synthesis via the SLC7A5–ERBB2–ACLY axis [40]. Altogether, these findings underscore that the resistance or lack of sensitivity to PARP inhibition is not driven by a single pathway but rather emerges from coordinated stress response and survival strategies in the context of DNA damage.

Despite these advances, the contribution of transcriptional regulators to cell survival via DNA damage response in BRCA2-mutated EOC remains poorly understood. In our study we have identified that KLF4 serves as a central transcriptional regulator that is activated by DNA damage in the context of BRCA2 mutations or HR deficiency in EOC. Because BRCA2 mutations or HR deficiency are deleterious to genomic stability, the inverse relationship between BRCA2 and KLF4 (**Fig 1**) suggests that up-regulation of KLF4-mediated signaling functions as a compensatory mechanism for HR deficiency in BRCA2-mutated EOC. Given the prominent role of KLF4 in stem cell reprogramming, DNA damage-induced KLF4 potentially increases the apoptotic threshold, conducing to increased cell survival and emerging PARP inhibitor resistance in BRCA2-mutated EOC.

Based on our bioinformatics analysis, we further identified that KLF4 mediates the up-regulation of genes including NR4A1 and p21 in BRCA2-mutated EOC cells (**Supplementary Table 1** and **Fig 3C**). Among these, NR4A1 emerged as the strongest KLF4-associated gene. NR4A1 is an immediate early-response transcription factor induced by a variety of physiological stresses [33]. NR4A1 displays diverse functions in cancer. Its overexpression has been linked to sarcoma progression [41] and worse outcome of high grade serous EOC [42]. p21 is a cyclin-dependent kinases inhibitor that blocks the G1/S transition in the cell cycle [43]. It is transcriptionally induced by p53 and KLF4 in response to DNA damage and stress conditions [44,45]. The loss of p21 has been demonstrated to convert KLF4 from tumor suppressing into tumor promoting properties [46]. Our study identified that KLF4 is required for the induction of both NR4A1 and p21 following PARP inhibition in BRCA2-mutated EOC cells (**Fig 4A**). It is plausible that p21 coordinates with the apoptotic program of p53 in response to DNA damage. With the majority of EOC harboring p53 mutations, we postulate that KLF4 dominates DNA damage responsive programs and favors survival in the absence of p53-mediated apoptosis. Indeed, it is supported by our results showing that silencing KLF4 or NR4A1, but not p21, increases PARP inhibition-induced apoptosis in BRCA2-mutated EOC cells (**Fig 4B**).

BH3 mimetic inhibitors have been evaluated in pre-clinical models and clinical trials for the treatment of EOC [47–49]. We found that navitoclax, a BH3 mimetic inhibitor targeting BCL2, BCL-xL, and BCL-w, selectively reduced survival and enhanced the apoptotic effects of olaparib on BRCA2-mutated EOC cells (**Fig 5A-C**). In contrast, other BH3 mimetic inhibitors sabutoclax, venetoclax, and the MCL1-specific A1210477 enhanced the efficacy of olaparib equally against BRCA2-mutated and BRCA2-wild type EOC cells (**Supplementary Fig 1** and **Supplementary Fig 2**), possibly due to lacking inhibitory activity against BCL-w (**Supplementary Table 2**). This finding was further supported by our results of bioinformatic analysis and western blotting, showing that KLF4 directly up-regulated BCL-xL and BCL-w and that KLF4 silencing abrogated olaparib-induced BCL-xL and BCL-w in BRCA2-mutated EOC cells (**Fig 6A, D**).

Unexpectedly, silencing of KLF4 increased the baseline levels of BCL-w in PEO1-NR cells and BCL-xL in both PEO1 and PEO1-NR cells. The baseline levels of NR4A1 and cIAP2 were noticeably elevated in PEO1-NR cells (**Fig 6B, E, F**). These observations suggest the existence of adaptive or compensatory mechanism acquired along with PARP inhibitor resistance in PEO1-NR cells. Providing that NR4A1 can directly up-regulate cIAP2 (**Fig 6A**) and silencing of KLF4 had minimal effects on olaparib-induced cIAP2 (**Fig 6E**), we speculate that NR4A1 can up-regulate cIAP2 by DNA damage independently of KLF4. This phenomenon is supported by the finding that birinapant, a SMAC inhibitor [50], enhanced olaparib-induced apoptosis in both BRCA2-mutated and BRCA2-wild type EOC cells (**Fig 5D, E**).

Taken together with our findings, we propose a mechanism by which KLF4 promotes apoptosis evasion through bifurcate signaling pathways: first, the DNA damage-inducible BCL2 family proteins, and second, the DNA damage-inducible NR4A1-cIAP2 axis in BRCA2-mutated EOC cells (**Fig 8**). The NR4A1–cIAP2 axis may be activated by DNA damage independently of KLF4 both in BRCA2-mutated and BRCA2-wild type EOC cells. Jointly these pathways enable EOC cells to mitigate PARP inhibitor– induced DNA damage response, highlighting apoptosis-targeting strategies as rational combination approaches in different settings of BRCA2 or HR status. Furthermore, these pathways may shed light on the mechanisms by which, in addition to BRCA2 reversion mutations, BRCA2-mutated EOC develops therapeutic resistance (e.g. PARP inhibitors and platinum drugs) in patients who experience relapses and subsequently succumb to the disease.

**Figure 8.**
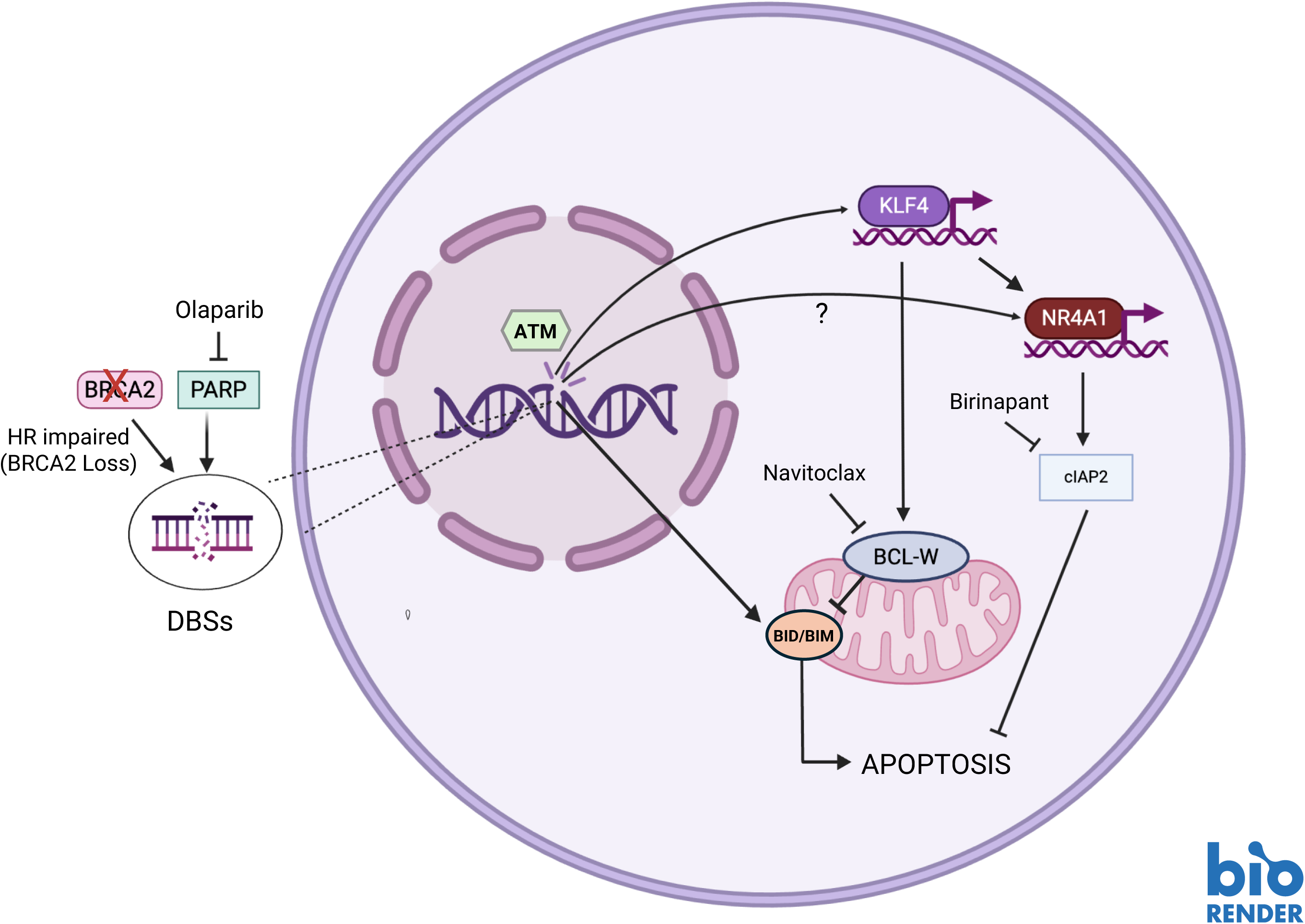
Proposed KLF4-driven survival signaling in BRCA2-mutated EOC. In these cells, treatment with the PARP inhibitor olaparib causes accumulation of unrepaired DSBs, leading to ATM-mediated KLF4 up-regulation. KLF4 promotes apoptosis evasion through two pathways: one being the induction of anti-apoptotic BCL-2 family proteins which suppress mitochondrial apoptosis by inhibiting BH3-only proteins (BID/BIM), and the other being the activation of the NR4A1–cIAP2 axis, which directly inhibits caspase-mediated apoptotic cascades. Pharmacological targeting of these pathways using navitoclax or birinapant restores apoptotic sensitivity. This proposed mechanism illustrates how KLF4-mediated signaling contributes to apoptosis evasion and therapeutic resistance in BRCA2-mutated EOC cells.

Our results from the tumor xenograft study further support the strategy that targeting anti-apoptotic proteins enhances the efficacy of PARP inhibitors and potentially prevents the development of PARP inhibitor-resistant in BRCA2-mutated EOC. In mice, navitoclax alone did not exhibit appreciable activity to deter EOC progression (**Fig 6**). Noticeably, it appeared to reduce the body weight of mice (about 9%) but show no other overt toxicity. We posit that non-selective inhibition of BCL2 family members by navitoclax may disrupt the homeostasis in normal tissues. It has been reported that navitoclax caused thrombocytopenia in a significant portion of patients due to inhibition of BCL-xL in platelets [51]. However, we observed that the combination of olaparib and navitoclax prominently reversed the reduction in body weight caused by navitoclax alone. Future development of BH3-mimetic inhibitors targeting BCL-w or BCL-xL will shed light on which BCL2 family member is responsible for anti-apoptotic activity in BRCA2-mutated EOC and address the issue of navitoclax-associated adverse effects. Navitoclax is not FDA-approved but it is encouraging that the Exactis-03 Phase I clinical trial (NCT05358639) is currently evaluating the safety and recommended dose of olaparib in combination with navitoclax for triple-negative breast cancer and high-grade serous EOC [52]. The study has not yet reported the efficacy outcomes.

In summary, we have identified the mechanism by which PARP inhibition induces KLF4 and consequently reduces apoptosis in BRCA2-mutated EOC cells. In addition, our study validates the strategy of combining olaparib and navitoclax to enhance the therapeutic efficacy for BRCA2-mutated EOC in patients. Despite the advances with PARP inhibitors, frequent recurrence and development of therapeutic resistance still pose a major challenge to successful treatment of high-grade serous EOC. Highly effective combination therapy based on targeting the vulnerability of EOC will make a transformative change in clinical practice and improve the survival outcome of EOC patients.

## Methods

### Cell Lines

Isogenic pairs of human BRCA2-mutated PEO1 and BRCA2-wild type PEO1-NR EOC cells, mouse p53(-/-);Brca2(+/+) (Brca2-WT) and p53(-/-);Brca2(-/-) (Brca2-KO) ID8 EOC cells were used for the experiments. PEO1 cells are patient-derived cell lines as described previously [34]. PEO1-NR cells were established by chronically exposing PEO1 cells to 2.5 μM niraparib for 4 weeks and propagating surviving niraparib-resistant PEO1 cells. All cell lines were cultured in DMEM medium supplemented with 10% FBS and 1% penicillin–streptomycin antibiotics. Brca2-WT and Brca2-KO ID8 cell lines were provided by Dr Iain McNeish from Imperial College London, UK [53].

### Chemicals and Antibodies

Olaparib (AZD2281) was purchased from SelleckChem. KU60019, ceralasertib, navitoclax (ABT-263), and birinapant were purchased from Cayman Chemicals. Human interferon-gamma (IFNγ) was purchased from Gemini Bio. Anti-γH2AX (Ser139) (RRID:AB_2118009), p-CHK1 (Ser345) (RRID:AB_331212), CHK1 (RRID: AB_2080320), KLF4 (RRID:AB_2797840), NR4A1 (RRID:AB_2153738), BCL2 (RRID:AB_2744528), BCL-xL (RRID:AB_2228008), BCL-w (RRID:AB_10691557), cIAP1 (RRID:AB_10890862), and cIAP2 (RRID:AB_10693298) antibodies were purchased from Cell Signaling. The anti-NR4A1 (RRID: AB_3750353) antibody was purchased from Boster. The anti-BCL-w (RRID:AB_10983656) was purchased from ThermoFisher. Anti-HSC70 (RRID:AB_627761) and β-Actin (RRID:AB_626632) antibodies were purchased from Santa Cruz. The anti-BRCA2 (RRID:AB_10952240) antibody was purchased from Bethyl/Fortis.

### DNA Sequencing

Genomic DNA was isolated from cells using the DNeasy Blood & Tissue kit (Qiagen). DNA sequencing primers spanning the region that contains the nonsense mutation (5193C>G) of the BRCA2 protein in PEO1 cells were synthesized by Yale Keck Oligo Synthesis Facility. They were: 5’ GCTGCCCCAAAGTGTAAAGA-3’ and 5’-GGTGAAGAGCTAGTCACAAGT-3’. DNA sequencing was performed using the Sanger method at Yale Keck DNA sequencing facility. The sequences were visualized using the Snapgene software.

### Western Blotting

The methodology was described previously [18]. All images were acquired and processed with the G:BOX gel documentation system and the GeneSnap software (Syngene).

### CRISPR Gene Editing

The oligos for the sgRNA sequence of BRCA2, 5’-CACCGGTCTACCTGACCAATCGATG-3’ and 5’-AAACCATCGATTGGTCAGGTAGACC-3’ were designed using the CRISPick website (https://portals.broadinstitute.org/gppx/crispick/public) and synthesized by Yale Keck Oligo Synthesis Facility. These oligos were annealed and ligated into BbstI-digested pSpCas9(BB)-2A-Puro V2.0 (Addgene; Plasmid #62988). PEO1-NR cells were transfected with the resulting BRCA2 gRNA-expressing construct and selected with puromycin. Puromycin-resistant clones were expanded to cell lines and screened for BRCA2 knockout using western blot analysis.

### Caspase 3/7 Assay

The methodology was described previously [34,54]. Following drug treatment, cells were lysed using the lysis buffer (PBS, 1% NP40, 0.1% SDS). Ten µL of lysate was incubated with Caspase-Glo 3/7 Assay reagent (Promega) for 1 hr at room temperature. Luminescence was detected by a luminometer. The protein concentration of lysate was measured with the DC Protein Assay (Bio-Rad) according to the manufacturer’s instructions. Caspase-3/7 activity was normalized to the amount of protein in the lysate and expressed as RLU per μg of protein.

### Quantitative RT-PCR

Total RNA was extracted from cells using RNeasy Kit (Qiagen). PCR primers were designed using NCBI Primer-Blast website and synthesized by Yale Keck Oligo Synthesis Facility. They were: KLF4, 5’AACACACCGGGTTAATAAGGCA-3’and 3’-GCAAACCTGAACTGGGTTTTT-5’; GAPDH, 5’-ACAGTCCATGCCATCACTGCC-3’ and 3’-GCCTGCTTCACCACCTTCTTG-5’; NR4A1, 5’-TGACTACTATGGCAGCCCCT-3’ and 3’-CCTTCGTAAGTCTGGCTGGG-5’; CDKN1A, 5’-TGGGCTTTTGGGAAGGATTTCA-3’and 3’ATCTTTGCAGAGTGTGACAGT-5’; mouse KLF4, 5’-CTTACAGCCTTCCGAGGTGG-3’ and 5’-GAAGACTAGTGGGGAACCGC-3’; mouse GAPDH, 5’-AATACGGCTACAGCAACAGGG-3’ and 5’-CTCTTGCTCAGTGTCCTTGC-3’. All PCR reactions were performed using the iTaq Universal One-Step RT-qPCR Kit (Bio-Rad) in the Mx3000p real-time PCR thermal cycler (Strategene): one cycle for 10 min at 50 °C and one cycle for 1 min at 95°C followed by 40 cycles 15 s at 95 °C and 30 s at 60 °C. Determination of fold gene expression was performed using the 2 ^ΔΔCt^ method.

### MTS Cytotoxicity/Viability Assay

The methodology was described previously [34,54]. Cells were plated into 96 well plates for 24 hours. Drugs in single or combinations were added in triplicate wells for 72 hours. Thereafter, 20 µL of CellTiter 96 AQueous MTS Reagent (Promega) was added to each well. After 2-hour incubation the plates were read at 490 nm by a colorimetric microplate reader. The percentage of cell survival was determined by calculating absorbance and normalizing the values to vehicle-treated controls.

### Gene Silencing

The methodology was detailed previous [18,55]. Cells were plated into 6 well plates, incubated for 24 h, and transfected with 50 nM siRNA using a Lipofectamine 2000 transfection reagent (ThermoFisher) according to the manufacturer’s protocol. Transfected cells were incubated for 48 hr before assays. All siRNAs were purchased from Millipore-Sigma. siRNA sequences are as follows: Non-target control (NTC), siRNA universal negative control#1 (SIC001). KLF4, 5’-GUGGAUAUCAGGGUAUAAAdTdT-3’. NR4A1, 5’-GGCUUGAGCUGCAGAAUGAdTdT-3’. CDKN1A, 5’-CUAAGAGUGCUGGGCAUUUdTdT-3’.

### Combination Synergy Analysis

Cell viability data were analyzed for the synergistic effect between two drugs using SynergyFinderPlus website (https://synergyfinderplus.org/) [56]. The average score of Loewe and ZIP methods and three-dimensional synergy surface plots was shown.

### Tumor Xenografts and Drug Treatment

The procedure was described previously [34]. The Yale University Institutional Animal Care and Use Committee approved the protocol for the in vivo animal studies in compliance with the US Public Health Policy on Humane Care and Use of Laboratory Animals. Five to six weeks old female SCID-Beige mice were obtained from Taconic Biosciences. PEO1ip cells suspended in 100 µl serum-free medium were implanted intraperitoneally (i.p.) (10 x 10^6^ cells per mouse). Mice were randomly assigned to treatment groups and treatment was initiated one day after implantation. Mice (n=5) received i.p. treatment with vehicle (DMSO), olaparib (50 mg/kg), navitoclax (50 mg/kg), or both drugs in combination once daily for 5 consecutive days per week for a total of 4 weeks. The body weight of mice was measured once a week for the first 7-8 weeks and every 1-3 days thereafter. The i.p. dissemination of PEO1ip xenografts mainly caused cachexia as detected by a progressive decline in body weight after 50 days [34]. The body condition score (BCS) of 2 [57], including hunched posture, reduced activity, and paleness, was defined as the survival endpoint. In some mice, development of tumor ascites (a sudden weight gain and increased abdominal circumference) with reduced activity was also defined as the endpoint. The Kaplan-Meier survival curve for each treatment group was generated, and the median survival time, statistical significance, and hazard ratio were determined using the Prism software (GraphPad).

## Statistical Analysis

Statistical analysis for cell-based assays was performed using unpaired t-test. All tests were two-tailed with an α level at 0.05.

### Bioinformatics Analysis

The cBioPortal for Cancer Genomics (https://www.cbioportal.org/)-hosting 6 datasets of total 1047 EOC patients were used for Kaplan-Meier survival analysis of KLF4 high and KLF4 low expression cohorts. The TCGA (GDC) dataset containing 602 high-grade serous EOC patients was used to identify genes that were highly correlated with KLF4 and NR4A1 in apoptosis, DNA damage response, and EMT/cancer stemness. The physical and functional associations in gene networks were established using the STRING database (https://string-db.org/). In addition, the edge weight of apoptotic gene network centered on KLF4 and NR4A1 in PEO1 cells was performed using the GRAND database. (https://grand.networkmedicine.org/networks/targeting/ACH_001630/).

## Acknowledgements

We thank Dr. Yasmin Korayem for her contribution to establishing the PEO1-NR cell line. This work was supported in part by grants from Rivkin Center for Ovarian Cancer Research (Z.P.L) and Peter E.Schwartz Discovery to Cure Program at Yale University (Z.P.L). Z Ping Lin is a Discovery to Cure Fellow.

## Author Contributions

Esma Fera: Conceptualization, Formal analysis, Investigation, Writing - Original Draft, Writing - Review & Editing, Visualization. Tiffany Zhang: Formal analysis, Investigation, Writing - Original Draft. Victoria M Grechukhina: Formal analysis, Investigation, Writing - Original Draft. Yong-Lian Zhu: Formal analysis, Investigation, Elena S Ratner: Resources, Writing - Review & Editing, Supervision, Project administration, Funding acquisition. Z Ping Lin: Conceptualization, Methodology, Validation, Formal analysis, Investigation, Resources, Writing - Original Draft, Writing - Review & Editing, Visualization, Funding acquisition.

## Data Availability

The data generated in this study are available within the article and its supplementary data files. Some of the data analyzed in this study were obtained from the cBioPortal for Cancer Genomics’ ovarian cancer datasets. Functional and protein interaction and gene regulatory network analyses were performed using publicly available resources, including the STRING database and the Gene Regulatory Network Database (GRAND).

## References

1 Society, A. C. Key Statistics for Ovarian Cancer, <https://www.cancer.org/cancer/types/ovarian-cancer/key-statistics.html#ovarian-cancer-estimates-for-2025> (2026).

2 Alsop, K. et al. BRCA mutation frequency and patterns of treatment response in BRCA mutation-positive women with ovarian cancer: a report from the Australian Ovarian Cancer Study Group. J Clin Oncol 30, 2654–2663, doi:10.1200/jco.2011.39.8545 (2012).

3 Ratner, E. S., Sartorelli, A. C. & Lin, Z. P. Poly (ADP-ribose) polymerase inhibitors: on the horizon of tailored and personalized therapies for epithelial ovarian cancer. Curr Opin Oncol 24, 564–571, doi:10.1097/CCO.0b013e3283564230 (2012).

4 Arora, S. et al. FDA Approval Summary: Olaparib Monotherapy or in Combination with Bevacizumab for the Maintenance Treatment of Patients with Advanced Ovarian Cancer. Oncologist 26, e164–e172, doi:10.1002/onco.13551 (2021).

5 Yoshida, T. in Encyclopedia of Signaling Molecules (ed Sangdun Choi) 2777–2781 (Springer International Publishing, 2018).

6 Shields, J. M., Christy, R. J. & Yang, V. W. Journal of Biological Chemistry 271, 20009–20017, doi:10.1074/jbc.271.33.20009 (1996).

7 Shields, J. M. & Yang, V. W. Two Potent Nuclear Localization Signals in the Gut-enriched Krüppel-like Factor Define a Subfamily of Closely Related Krüppel Proteins*. Journal of Biological Chemistry 272, 18504–18507, 10.1074/jbc.272.29.18504 (1997).

8 Takahashi, K. & Yamanaka, S. Induction of pluripotent stem cells from mouse embryonic and adult fibroblast cultures by defined factors. Cell 126, 663–676, doi:10.1016/j.cell.2006.07.024 (2006).

9 Aguirre, M. et al. Application of the Yamanaka Transcription Factors Oct4, Sox2, Klf4, and c-Myc from the Laboratory to the Clinic. Genes 14, 1697 (2023).

10 Rowland, B. D., Bernards, R. & Peeper, D. S. The KLF4 tumour suppressor is a transcriptional repressor of p53 that acts as a context-dependent oncogene. Nature Cell Biology 7, 1074–1082, doi:10.1038/ncb1314 (2005).

11 Wang, X. et al. The deubiquitinase USP10 regulates KLF4 stability and suppresses lung tumorigenesis. Cell Death & Differentiation 27, 1747–1764, doi:10.1038/s41418-019-0458-7 (2020).

12 Song, X. et al. Expression of Krüppel-like factor 4 in breast cancer tissues and its effects on the proliferation of breast cancer MDA-MB-231 cells. Exp Ther Med 13, 2463–2467, doi:10.3892/etm.2017.4262 (2017).

13 Wang, B. et al. Krüppel-like factor 4 induces apoptosis and inhibits tumorigenic progression in SK-BR-3 breast cancer cells. FEBS Open Bio 5, 147–154, doi:10.1016/j.fob.2015.02.003 (2015).

14 Yu, F. et al. Kruppel-like factor 4 (KLF4) is required for maintenance of breast cancer stem cells and for cell migration and invasion. Oncogene 30, 2161–2172, doi:10.1038/onc.2010.591 (2011).

15 Ghaleb, A. M., Katz, J. P., Kaestner, K. H., Du, J. X. & Yang, V. W. Krüppel-like factor 4 exhibits antiapoptotic activity following gamma-radiation-induced DNA damage. Oncogene 26, 2365– 2373, doi:10.1038/sj.onc.1210022 (2007).

16 Zhou, W. et al. KLF4 promotes cisplatin resistance by activating mTORC1 signaling in ovarian cancer. Discov Oncol 15, 682, doi:10.1007/s12672-024-01576-y (2024).

17 Wang, B. et al. KLF4 expression enhances the efficacy of chemotherapy drugs in ovarian cancer cells. Biochem Biophys Res Commun 484, 486–492, doi:10.1016/j.bbrc.2017.01.062 (2017).

18 Lin, A. C., Moscarelli, J., Zhu, Y.-L., Lin, Z. P. & Ratner, E. S. CXCL10-induced regulatory T cells and adenosine signaling promote immunosuppression and progression of epithelial ovarian cancer. Scientific Reports 15, 20778, doi:10.1038/s41598-025-06812-1 (2025).

19 Wei, D., Kanai, M., Huang, S. & Xie, K. Emerging role of KLF4 in human gastrointestinal cancer. Carcinogenesis 27, 23–31, doi:10.1093/carcin/bgi243 (2006).

20 Blackford, A. N. & Jackson, S. P. ATM, ATR, and DNA-PK: The Trinity at the Heart of the DNA Damage Response. Molecular Cell 66, 801–817, doi:10.1016/j.molcel.2017.05.015 (2017).

21 Golding, S. E. et al. Improved ATM kinase inhibitor KU-60019 radiosensitizes glioma cells, compromises insulin, AKT and ERK prosurvival signaling, and inhibits migration and invasion. Mol Cancer Ther 8, 2894–2902, doi:10.1158/1535-7163.Mct-09-0519 (2009).

22 Vendetti, F. P. et al. The orally active and bioavailable ATR kinase inhibitor AZD6738 potentiates the anti-tumor effects of cisplatin to resolve ATM-deficient non-small cell lung cancer in vivo. Oncotarget 6, 44289–44305, doi:10.18632/oncotarget.6247 (2015).

23 Yoon, H. S., Chen, X. & Yang, V. W. Kruppel-like factor 4 mediates p53-dependent G1/S cell cycle arrest in response to DNA damage. J Biol Chem 278, 2101–2105, doi:10.1074/jbc.M211027200 (2003).

24 Farrugia, M. K., Vanderbilt, D. B., Salkeni, M. A. & Ruppert, J. M. Kruppel-like Pluripotency Factors as Modulators of Cancer Cell Therapeutic Responses. Cancer Res 76, 1677–1682, doi:10.1158/0008-5472.can-15-1806 (2016).

25 Szklarczyk, D. et al. The STRING database in 2023: protein-protein association networks and functional enrichment analyses for any sequenced genome of interest. Nucleic Acids Res 51, D638–d646, doi:10.1093/nar/gkac1000 (2023).

26 Youle, R. J. & Strasser, A. The BCL-2 protein family: opposing activities that mediate cell death. Nature Reviews Molecular Cell Biology 9, 47–59, doi:10.1038/nrm2308 (2008).

27 Diepstraten, S. T. et al. The manipulation of apoptosis for cancer therapy using BH3-mimetic drugs. Nature Reviews Cancer 22, 45–64, doi:10.1038/s41568-021-00407-4 (2022).

28 Tse, C. et al. ABT-263: a potent and orally bioavailable Bcl-2 family inhibitor. Cancer Res 68, 3421–3428, doi:10.1158/0008-5472.Can-07-5836 (2008).

29 Wei, J. et al. BI-97C1, an optically pure Apogossypol derivative as pan-active inhibitor of antiapoptotic B-cell lymphoma/leukemia-2 (Bcl-2) family proteins. J Med Chem 53, 4166–4176, doi:10.1021/jm1001265 (2010).

30 Souers, A. J. et al. ABT-199, a potent and selective BCL-2 inhibitor, achieves antitumor activity while sparing platelets. Nat Med 19, 202–208, doi:10.1038/nm.3048 (2013).

31 Leverson, J. D. et al. Potent and selective small-molecule MCL-1 inhibitors demonstrate on-target cancer cell killing activity as single agents and in combination with ABT-263 (navitoclax). Cell Death Dis 6, e1590, doi:10.1038/cddis.2014.561 (2015).

32 Krepler, C. et al. The Novel SMAC Mimetic Birinapant Exhibits Potent Activity against Human Melanoma Cells. Clinical Cancer Research 19, 1784–1794, doi:10.1158/1078-0432.CCR-12-2518 (2013).

33 Maxwell, M. A. & Muscat, G. E. The NR4A subgroup: immediate early response genes with pleiotropic physiological roles. Nucl Recept Signal 4, e002, doi:10.1621/nrs.04002 (2006).

34 Lin, Z. P. et al. Combination of triapine, olaparib, and cediranib suppresses progression of BRCA-wild type and PARP inhibitor-resistant epithelial ovarian cancer. PLOS ONE 13, e0207399, doi:10.1371/journal.pone.0207399 (2018).

35 Rose, P. G. et al. PARP inhibitors decrease response to subsequent platinum-based chemotherapy in patients with BRCA mutated ovarian cancer. Anticancer Drugs 32, 1086–1092, doi:10.1097/cad.0000000000001219 (2021).

36 Biegała, Ł., Gajek, A., Marczak, A. & Rogalska, A. Olaparib-Resistant BRCA2MUT Ovarian Cancer Cells with Restored BRCA2 Abrogate Olaparib-Induced DNA Damage and G2/M Arrest Controlled by the ATR/CHK1 Pathway for Survival. Cells 12, 1038 (2023).

37 Decker, J. T. et al. Systems analysis of dynamic transcription factor activity identifies targets for treatment in Olaparib resistant cancer cells. Biotechnology and Bioengineering 114, 2085–2095, 10.1002/bit.26293 (2017).

38 Norquist, B. et al. Secondary somatic mutations restoring BRCA1/2 predict chemotherapy resistance in hereditary ovarian carcinomas. J Clin Oncol 29, 3008–3015, doi:10.1200/jco.2010.34.2980 (2011).

39 Vaidyanathan, A. et al. ABCB1 (MDR1) induction defines a common resistance mechanism in paclitaxel-and olaparib-resistant ovarian cancer cells. British Journal of Cancer 115, 431–441, doi:10.1038/bjc.2016.203 (2016).

40 Liu, S. et al. SLC7A5-ERBB2 axis drives olaparib resistance via de novo lipid synthesis in ovarian cancer. Oncogene 45, 140–263, doi:10.1038/s41388-025-03584-w (2026).

41 Lacey, A., Rodrigues-Hoffman, A. & Safe, S. PAX3-FOXO1A Expression in Rhabdomyosarcoma Is Driven by the Targetable Nuclear Receptor NR4A1. Cancer Res 77, 732–741, doi:10.1158/0008-5472.Can-16-1546 (2017).

42 Delgado, E. et al. High expression of orphan nuclear receptor NR4A1 in a subset of ovarian tumors with worse outcome. Gynecol Oncol 141, 348–356, doi:10.1016/j.ygyno.2016.02.030 (2016).

43 Lin, Z. P., Zhu, Y. L. & Ratner, E. S. Targeting Cyclin-Dependent Kinases for Treatment of Gynecologic Cancers. Front Oncol 8, 303, doi:10.3389/fonc.2018.00303 (2018).

44 Zhang, W. et al. The gut-enriched Kruppel-like factor (Kruppel-like factor 4) mediates the transactivating effect of p53 on the p21WAF1/Cip1 promoter. J Biol Chem 275, 18391–18398, doi:10.1074/jbc.C000062200 (2000).

45 Karimian, A., Ahmadi, Y. & Yousefi, B. Multiple functions of p21 in cell cycle, apoptosis and transcriptional regulation after DNA damage. DNA Repair (Amst) 42, 63–71, doi:10.1016/j.dnarep.2016.04.008 (2016).

46 Rowland, B. D. & Peeper, D. S. KLF4, p21 and context-dependent opposing forces in cancer. Nat Rev Cancer 6, 11–23, doi:10.1038/nrc1780 (2006).

47 Dai, Y., Jin, S., Li, X. & Wang, D. The involvement of Bcl-2 family proteins in AKT-regulated cell survival in cisplatin resistant epithelial ovarian cancer. Oncotarget 8, 1354–1368, doi:10.18632/oncotarget.13817 (2017).

48 Del Bufalo, D. & Damia, G. Overview of BH3 mimetics in ovarian cancer. Cancer Treatment Reviews 129, 102771, 10.1016/j.ctrv.2024.102771 (2024).

49 Simonin, K. et al. Platinum compounds sensitize ovarian carcinoma cells to ABT-737 by modulation of the Mcl-1/Noxa axis. Apoptosis 18, 492–508, doi:10.1007/s10495-012-0799-x (2013).

50 Bai, L., Smith, D. C. & Wang, S. Small-molecule SMAC mimetics as new cancer therapeutics. Pharmacol Ther 144, 82–95, doi:10.1016/j.pharmthera.2014.05.007 (2014).

51 Rudin, C. M. et al. Phase II study of single-agent navitoclax (ABT-263) and biomarker correlates in patients with relapsed small cell lung cancer. Clin Cancer Res 18, 3163–3169, doi:10.1158/1078-0432.Ccr-11-3090 (2012).

52 Mackay, H. et al. Exactis-03: A phase I trial of the combination of olaparib and navitoclax in women with high grade serous ovarian cancer and triple negative breast cancer. Journal of Clinical Oncology 41, TPS5623–TPS5623, doi:10.1200/JCO.2023.41.16_suppl.TPS5623 (2023).

53 Walton, J. et al. CRISPR/Cas9-Mediated Trp53 and Brca2 Knockout to Generate Improved Murine Models of Ovarian High-Grade Serous Carcinoma. Cancer Research 76, 6118–6129, doi:10.1158/0008-5472.CAN-16-1272 (2016).

54 Lin, Z. P. et al. In silico screening identifies a novel small molecule inhibitor that counteracts PARP inhibitor resistance in ovarian cancer. Sci Rep 11, 8042, doi:10.1038/s41598-021-87325-5 (2021).

55 Lin, Z. P., Ratner, E. S., Whicker, M. E., Lee, Y. & Sartorelli, A. C. Triapine disrupts CtIP-mediated homologous recombination repair and sensitizes ovarian cancer cells to PARP and topoisomerase inhibitors. Mol Cancer Res 12, 381–393, doi:10.1158/1541-7786.Mcr-13-0480 (2014).

56 Yadav, B., Wennerberg, K., Aittokallio, T. & Tang, J. Searching for Drug Synergy in Complex Dose-Response Landscapes Using an Interaction Potency Model. Comput Struct Biotechnol J 13, 504–513, doi:10.1016/j.csbj.2015.09.001 (2015).

57 Ullman-Culleré, M. H. & Foltz, C. J. Body condition scoring: a rapid and accurate method for assessing health status in mice. Lab Anim Sci 49, 319–323 (1999).

