## Supplementary Tables and Figures for "KLF4 promotes apoptosis evasion and PARP inhibitor resistance in BRCA2-mutated epithelial ovarian cancer"

| Correlated Gene | Cytoband | Spearman's Correlation | p-Value | q-Value |
|---|---|---|---|---|
| NR4A1 | 12q13.13 | 0.6069 | 2.57E-44 | 1.01E-39 |
| SLC25A25 | 9q34.11 | 0.5792 | 1.26E-39 | 2.47E-35 |
| NR4A3 | 9q22 | 0.5677 | 8.49E-38 | 1.11E-33 |
| DUSP1 | 5q35.1 | 0.5646 | 2.54E-37 | 2.49E-33 |
| KLF10 | 8q22.3 | 0.5565 | 4.39E-36 | 3.44E-32 |
| CDKN1A | 6p21.2 | 0.5530 | 1.44E-35 | 9.41E-32 |
| THBD | 20p11.21 | 0.5431 | 3.99E-34 | 2.24E-30 |
| FOS | 14q24.3 | 0.5350 | 5.50E-33 | 2.48E-29 |
| ZFP36 | 19q13.2 | 0.5349 | 5.69E-33 | 2.48E-29 |
| FOSB | 19q13.32 | 0.5343 | 6.76E-33 | 2.65E-29 |
| EGR3 | 8p21.3 | 0.5328 | 1.12E-32 | 4.00E-29 |
| CSRNP1 | 3p22.2 | 0.5231 | 2.33E-31 | 7.63E-28 |
| ATF3 | 1q32.3 | 0.5184 | 9.73E-31 | 2.94E-27 |
| EGR1 | 5q31.2 | 0.5135 | 4.22E-30 | 1.18E-26 |
| EGR2 | 10q21.3 | 0.5120 | 6.56E-30 | 1.72E-26 |
| RHOB | 2p24.1 | 0.5116 | 7.34E-30 | 1.80E-26 |
| C11ORF96 | 11p11.2 | 0.5081 | 2.08E-29 | 4.81E-26 |
| DUSP5 | 10q25.2 | 0.5069 | 2.95E-29 | 6.42E-26 |
| PIM1 | 6p21.2 | 0.5063 | 3.54E-29 | 7.30E-26 |
| ETS2 | 21q22.2 | 0.5060 | 3.82E-29 | 7.49E-26 |
| LRRC8A | 9q34.11 | 0.5041 | 6.72E-29 | 1.25E-25 |
| PPP1R15A | 19q13.2 | 0.4986 | 3.22E-28 | 5.73E-25 |
| NFIL3 | 9q22.31 | 0.4876 | 6.86E-27 | 1.17E-23 |
| MCL1 | 1q21.2 | 0.4835 | 2.12E-26 | 3.46E-23 |
| TIPARP | 3q25.31 | 0.4798 | 5.72E-26 | 8.97E-23 |

**Supplementary Table 1.** Gene expression highly correlates with KLF4 in the dataset of high-grade serous ovarian cancer (TCGA, GDC). Top 25 genes correlative with KLF4 are ranked based on Spearman's correlation, p-value, and q-value.

| | BCL2 | BCL-xL | BCL-w | MCL1 |
|---|---|---|---|---|
| **Navitoclax** | + | + | + | |
| **Sabutoclax** | + | + | | + |
| **Venetoclax** | + | | | |
| **A1210477** | | | | + |

**Supplementary Table 2.** Specificities of BCL2 inhibitors. "+" indicates the reported inhibitory effect on individual BCL2 members tested in the cited references.

| Gene | Cytoband | μ in (A) KLF4 High | μ in (B) KLF4 Low | σ in (A) KLF4 High | σ in (B) KLF4 Low | Log2 Ratio | p-Value | q-Value | Higher expression in |
|---|---|---|---|---|---|---|---|---|---|
| **BCL2L2** | 14q11.2 | 4.17 | 3.86 | 0.61 | 0.61 | 0.30 | 4.97e-7 | 1.150e-5 | (A) KLF4 High |
| **BCL2L11** | 2q13 | 4.42 | 4.11 | 0.69 | 0.64 | 0.30 | 3.254e-6 | 5.068e-5 | (A) KLF4 High |
| **BCL2L1-AS1** | 20q11.21 | 0.77 | 0.59 | 0.58 | 0.53 | 0.18 | 8.258e-4 | 4.586e-3 | (A) KLF4 High |
| **BCL2L15** | 1p13.2 | 0.92 | 0.70 | 0.89 | 0.68 | 0.22 | 5.060e-3 | 0.0204 | (A) KLF4 High |
| **BCL2L13** | 22q11.21 | 4.13 | 3.99 | 0.51 | 0.56 | 0.13 | 9.499e-3 | 0.0337 | (A) KLF4 High |
| **BCL2** | 18q21.33 | 1.81 | 1.67 | 0.82 | 0.86 | 0.14 | 0.0834 | 0.189 | (A) KLF4 High |
| **BCL2L2-PABPN1** | 14q11.2 | 0.18 | 0.15 | 0.15 | 0.18 | 0.03 | 0.0952 | 0.208 | (A) KLF4 High |
| **BCL2L1** | 20q11.21 | 6.61 | 6.53 | 0.59 | 0.52 | 0.08 | 0.132 | 0.266 | (A) KLF4 High |
| **BCL2A1** | 15q25.1 | 3.09 | 2.94 | 1.13 | 1.13 | 0.14 | 0.191 | 0.345 | (A) KLF4 High |
| **BCL2L10** | 15q21.2 | 1.66 | 1.57 | 0.93 | 0.95 | 0.09 | 0.316 | 0.484 | (A) KLF4 High |
| **BCL2L14** | 12p13.2 | 1.33 | 1.26 | 0.75 | 0.70 | 0.07 | 0.333 | 0.500 | (A) KLF4 High |
| **BCL2L12** | 19q13.33 | 5.11 | 5.07 | 0.63 | 0.66 | 0.04 | 0.480 | 0.639 | (A) KLF4 High |

**Supplementary Table 3.** Comparison of BCL2 family member expression levels between KLF4 high and KLF4 low cohorts in high-grade serous EOC patients from the TCGA (GDC) dataset (N=602). BCL-w (BCL2L2, marked in red) ranks first based on the statistical significance (p and q values). The data are derived from the volcano plot in Fig 3D.

**A**

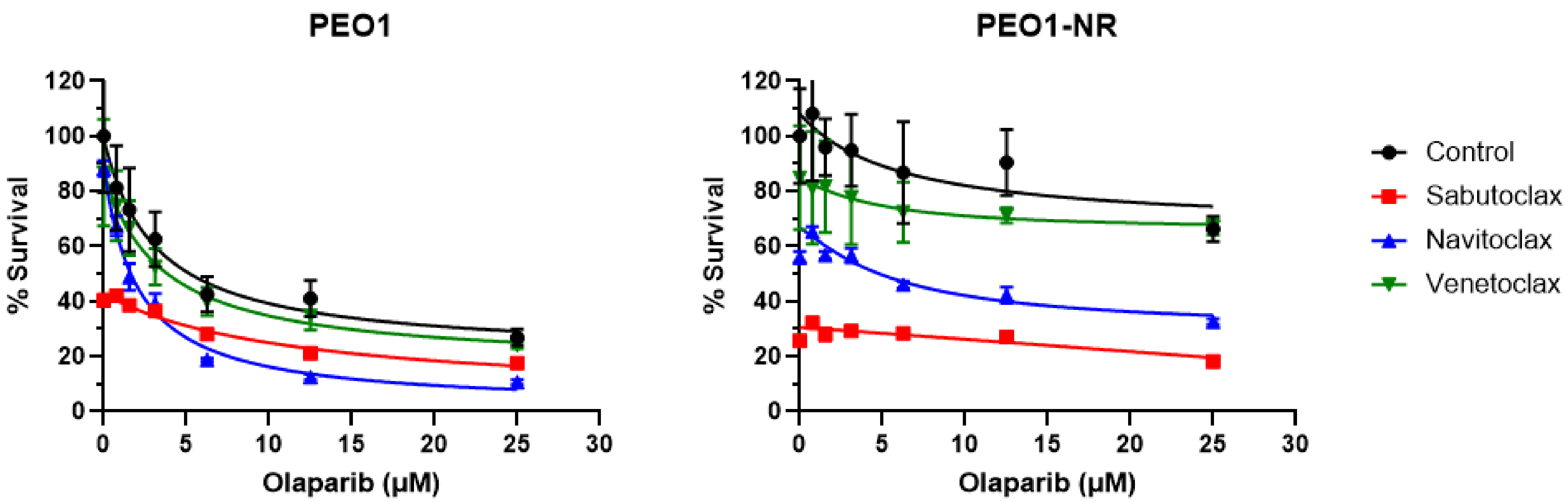


**Supplementary Figure 1.** The effects of BH3-mimetic inhibitors on the sensitivity of PEO1 and PEO1-NR cells to olaparib. **A.** PEO1 cells and PEO1-NR cells were treated with 6 μM sabutoclax, navitoclax, or venetoclax in combination with a range of concentrations of olaparib for 72 hr. Cell viability was determined by the MTS cytotoxicity assay and expressed as % survival. Data are means ± SD (n=3-4). **B.** Data of cell viability were used to determine the synergy score. Synergy scores using Loewe and ZIP methods are shown to evaluate the synergistic effect of the combination of olaparib and each BH3-mimetic inhibitor. Larger scores indicate greater synergistic effects between two drugs.

B

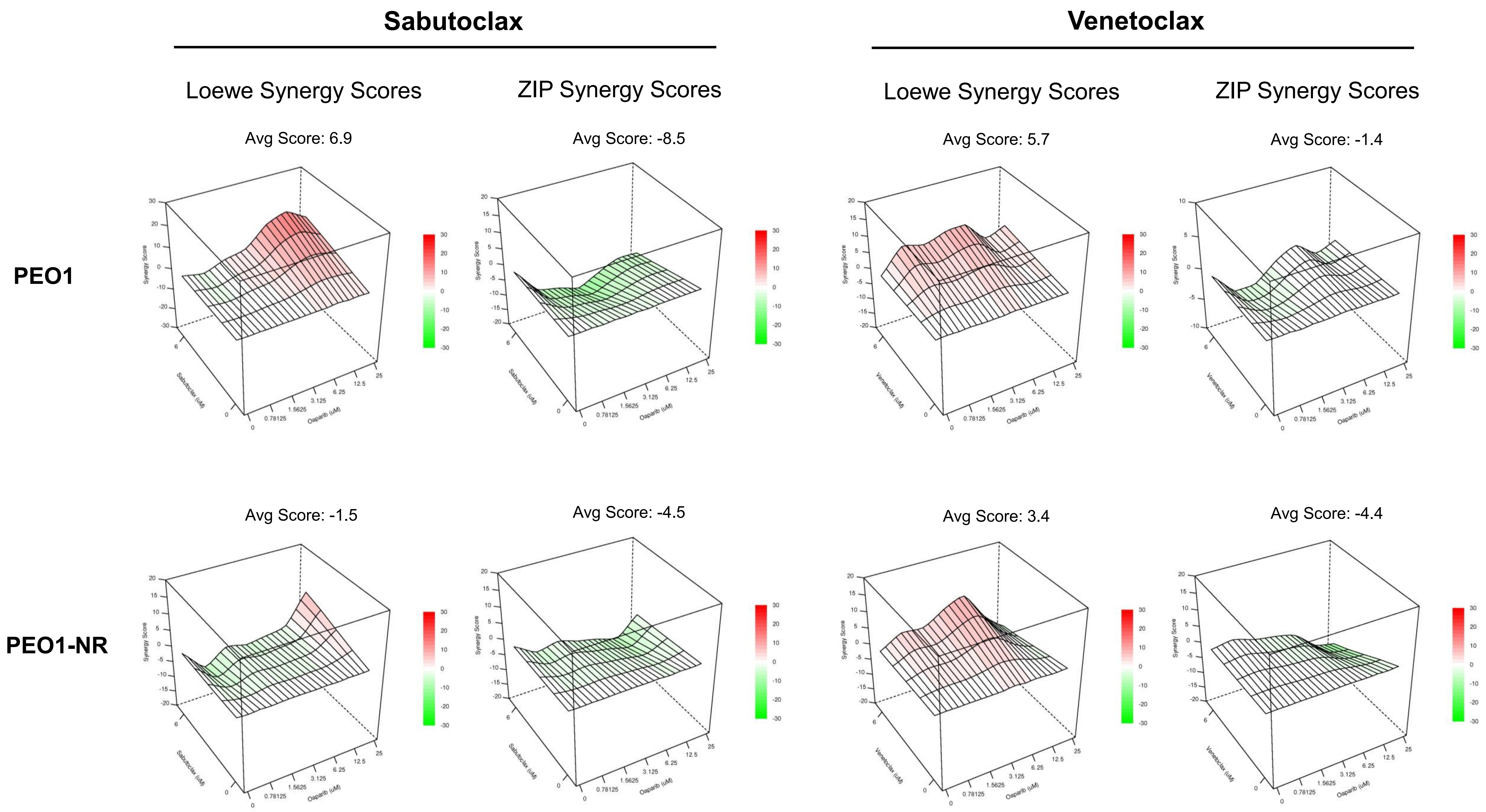


**Supplementary Figure 1-cont.**

**A**

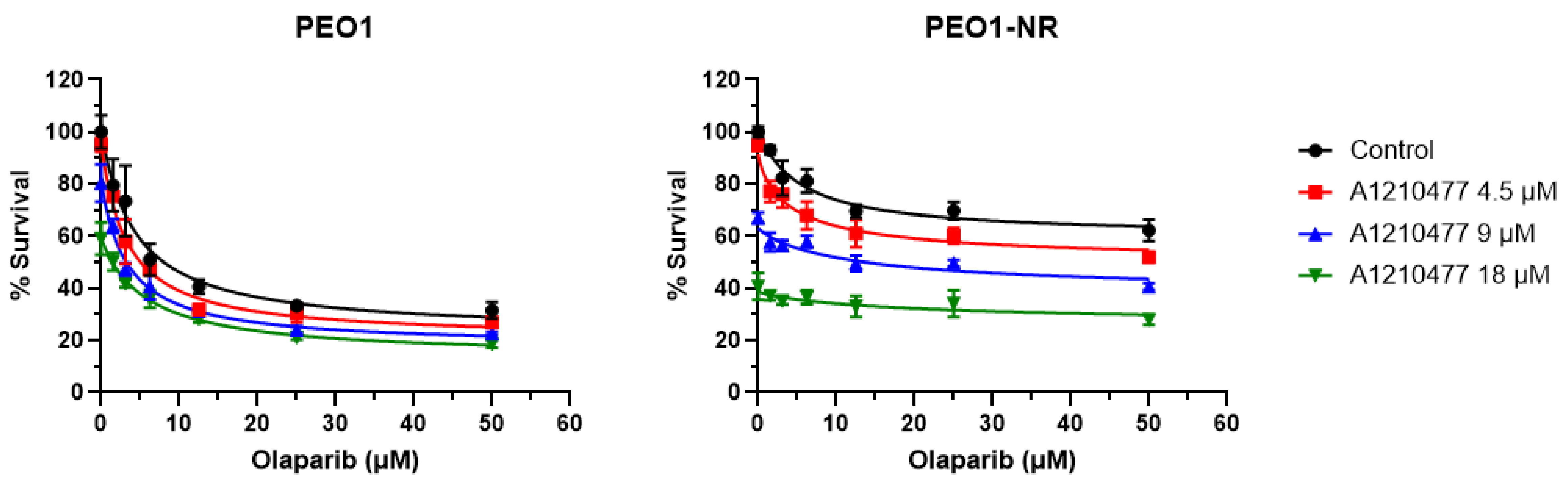


**Supplementary Figure 2.** The effects of A1210477 on the sensitivity of PEO1 and PEO1-NR cells to olaparib. **A.** PEO1 cells and PEO1-NR cells were treated with 4.5, 9, or 18 µM A1210477 in combination of a range of concentrations of olaparib for 72 hr. Cell viability was determined by the MTS cytotoxicity assay and expressed as % survival. **B.** Data of cell viability were used to determine the synergy score. Synergy scores using Loewe and ZIP methods are shown to evaluate the synergistic effect of the combination of olaparib and A1210477. Larger scores indicate greater synergistic effects between two drugs.

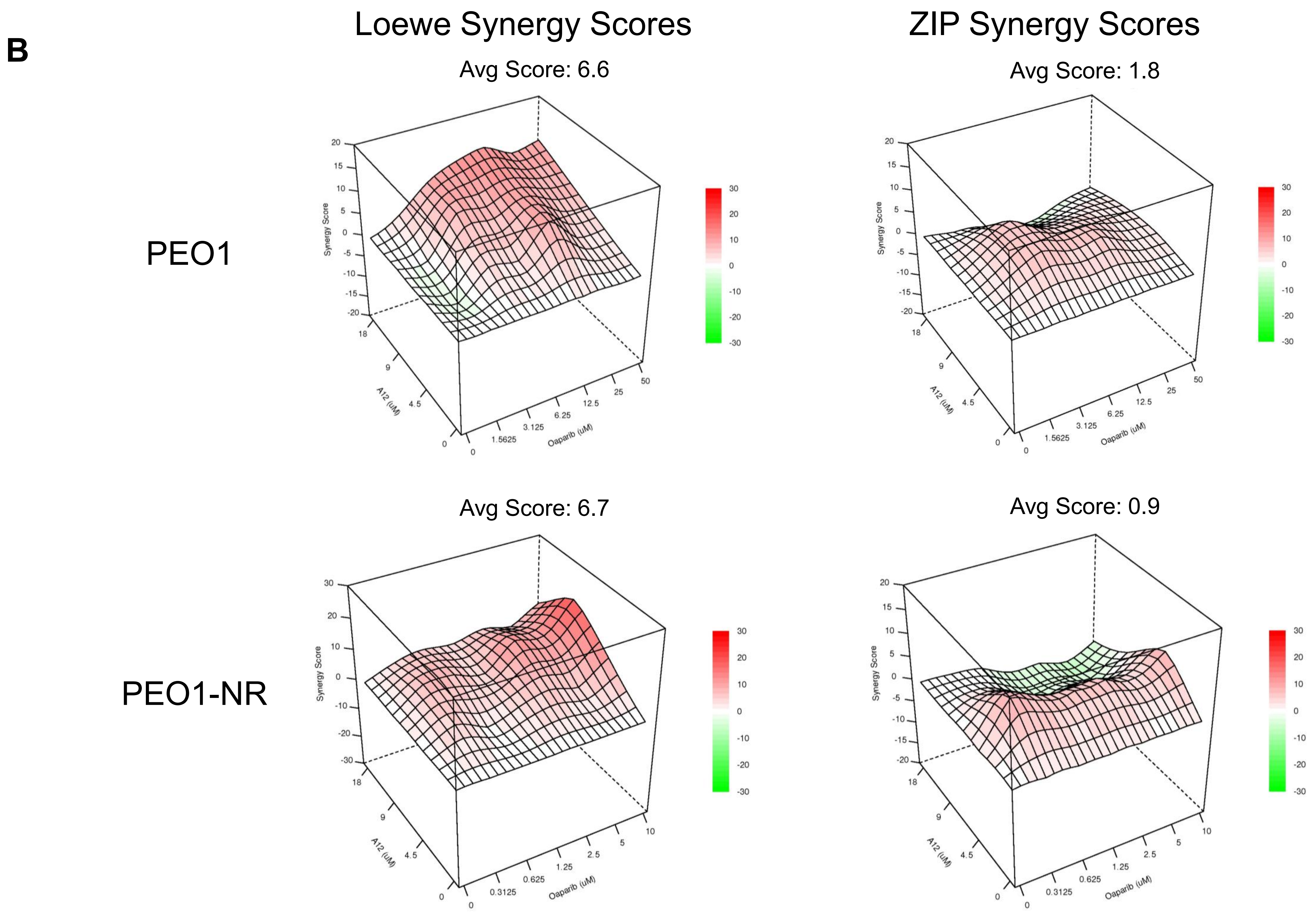


**Supplementary Figure 2-cont.**

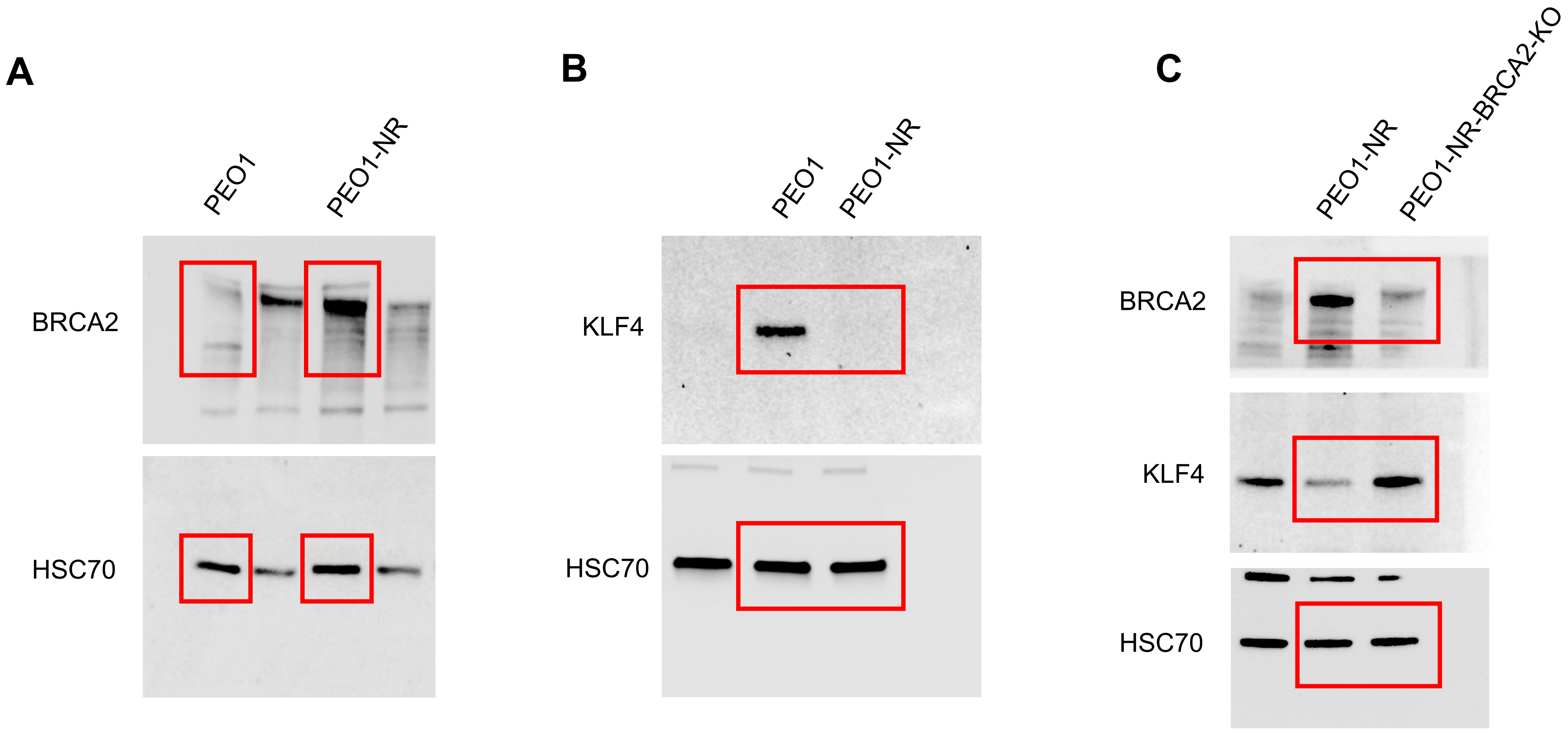


**Supplementary Figure 3A, B,C**, Images of uncropped blots for Figure 1B, C, E, respectively. Red outlines indicate cropped lanes.

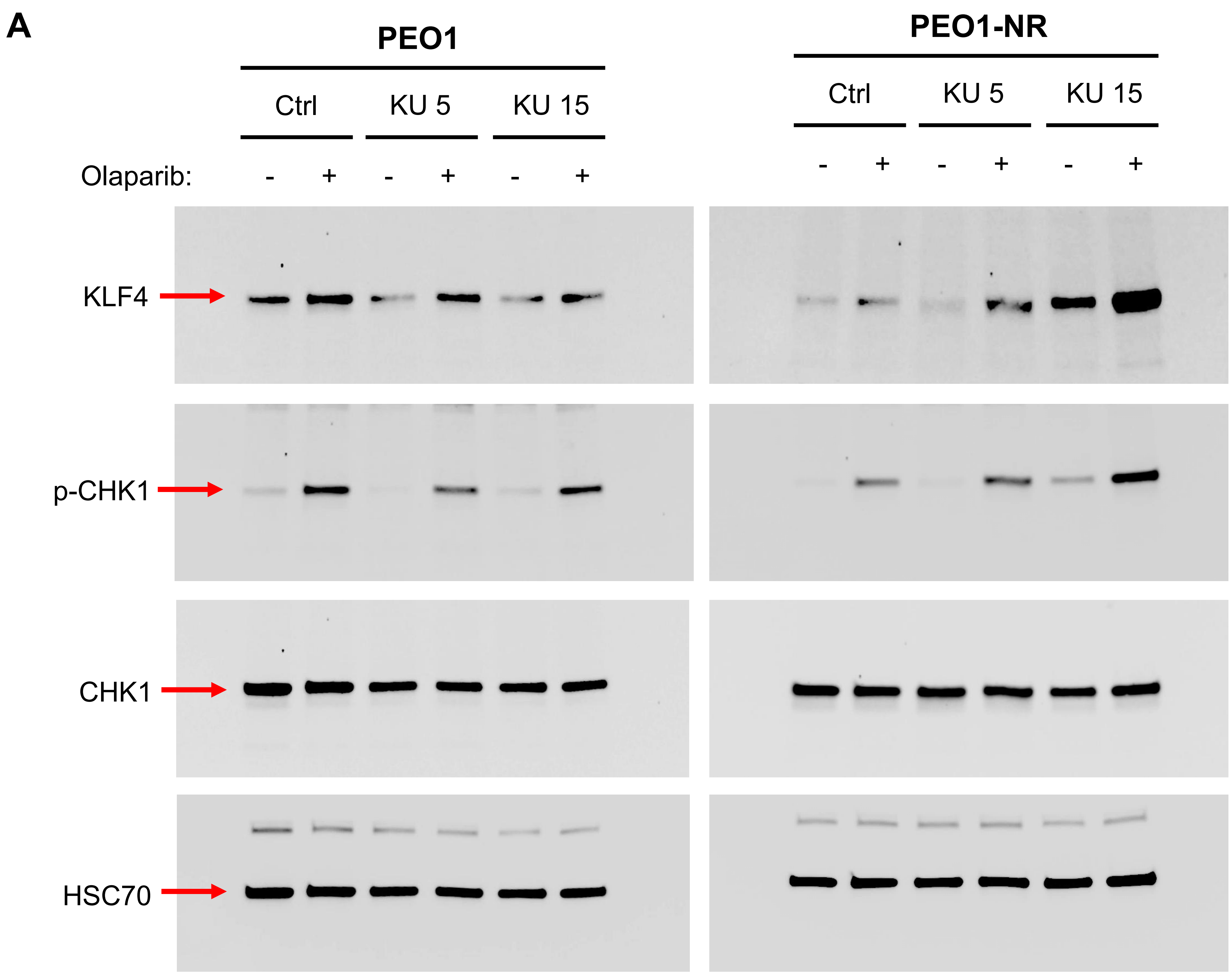


**Supplementary Figure 4A, B,** Images of uncropped blots for Figure 2, C, D, respectively.

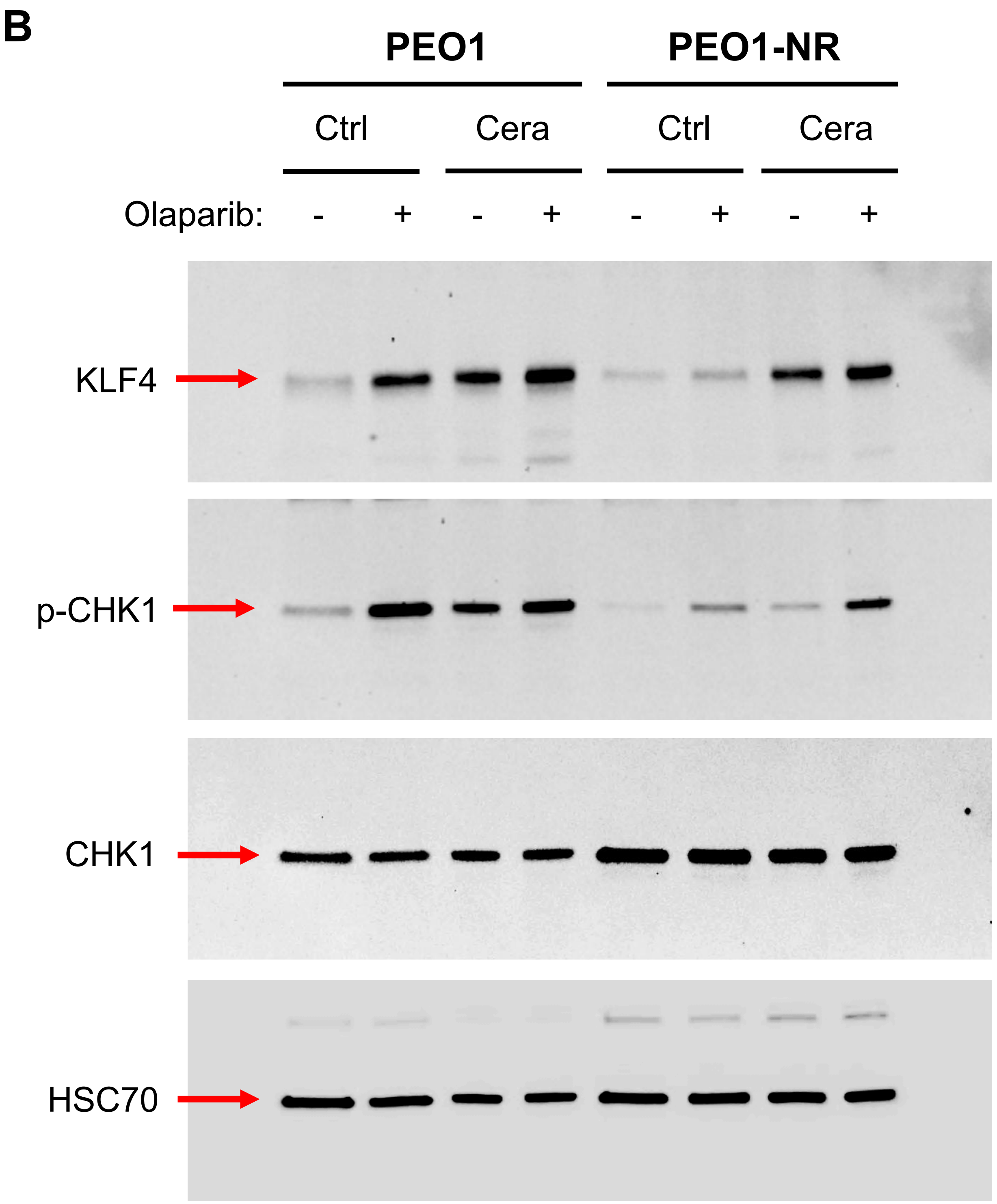


**Supplementary Figure 4-cont.**

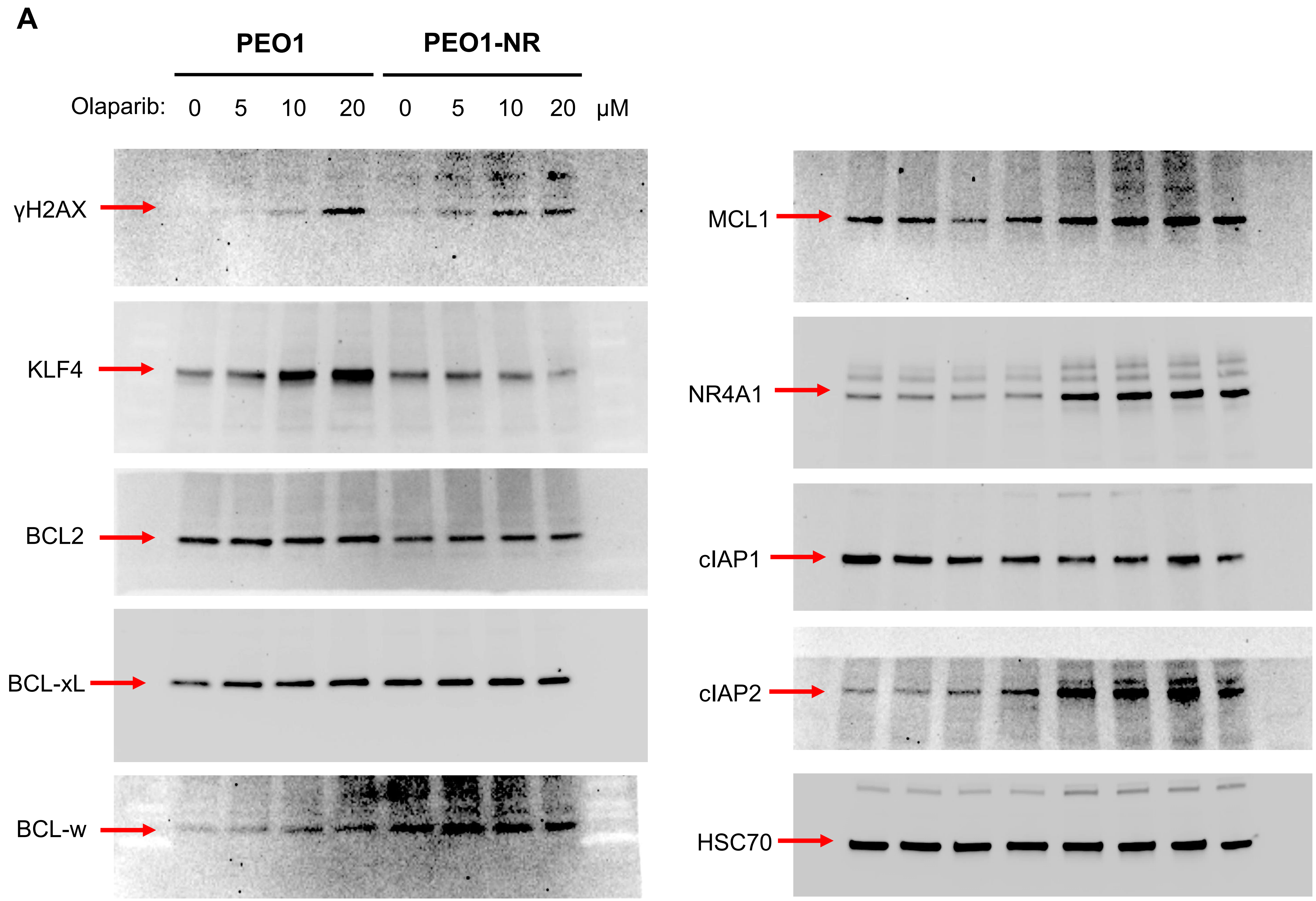


**Supplementary Figure 5A, B, C, D, E**, Images of uncropped blots for Figure 6B, C, D, E, F, respectively.

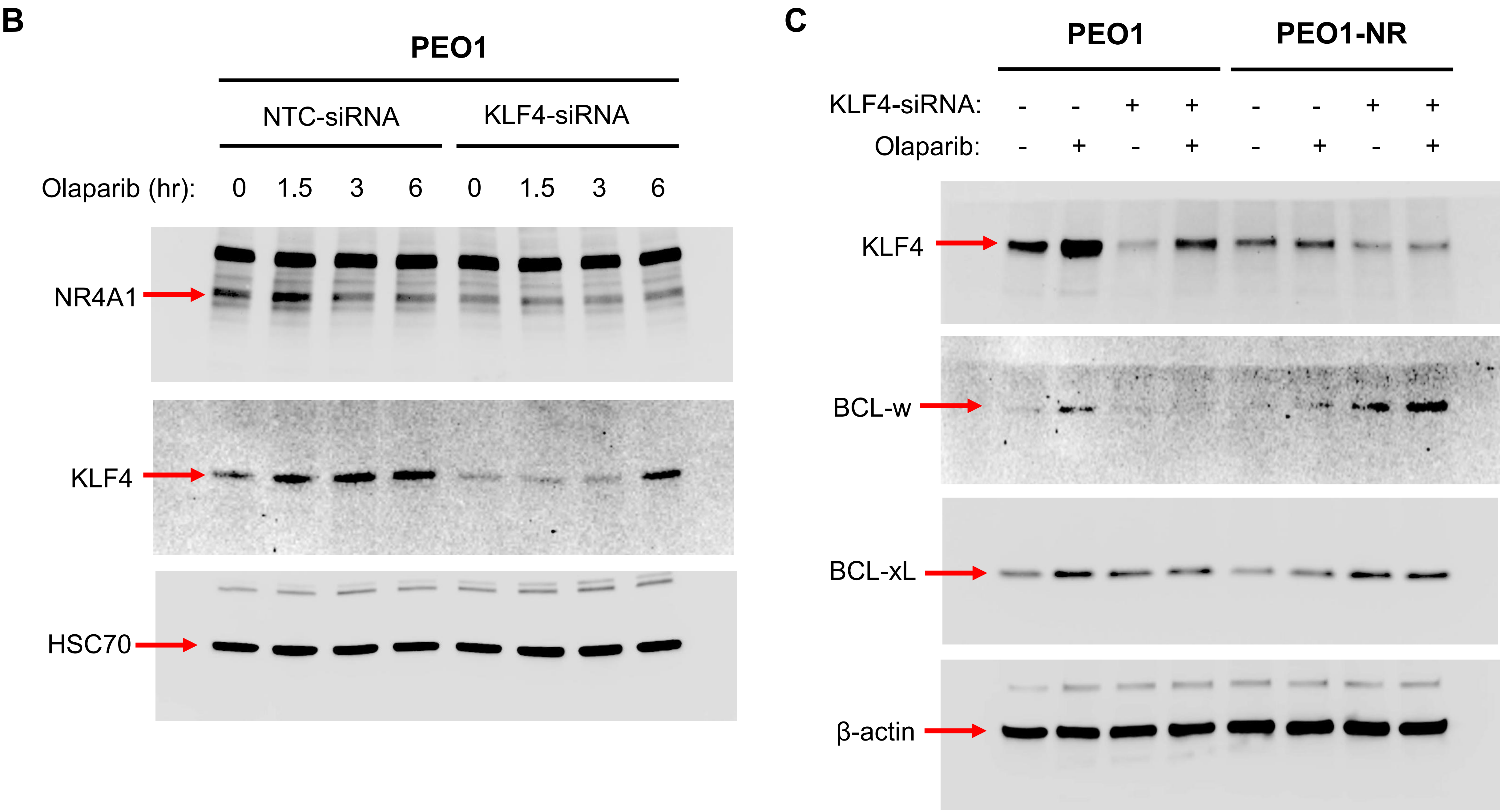


**Supplementary Figure 5-cont.**

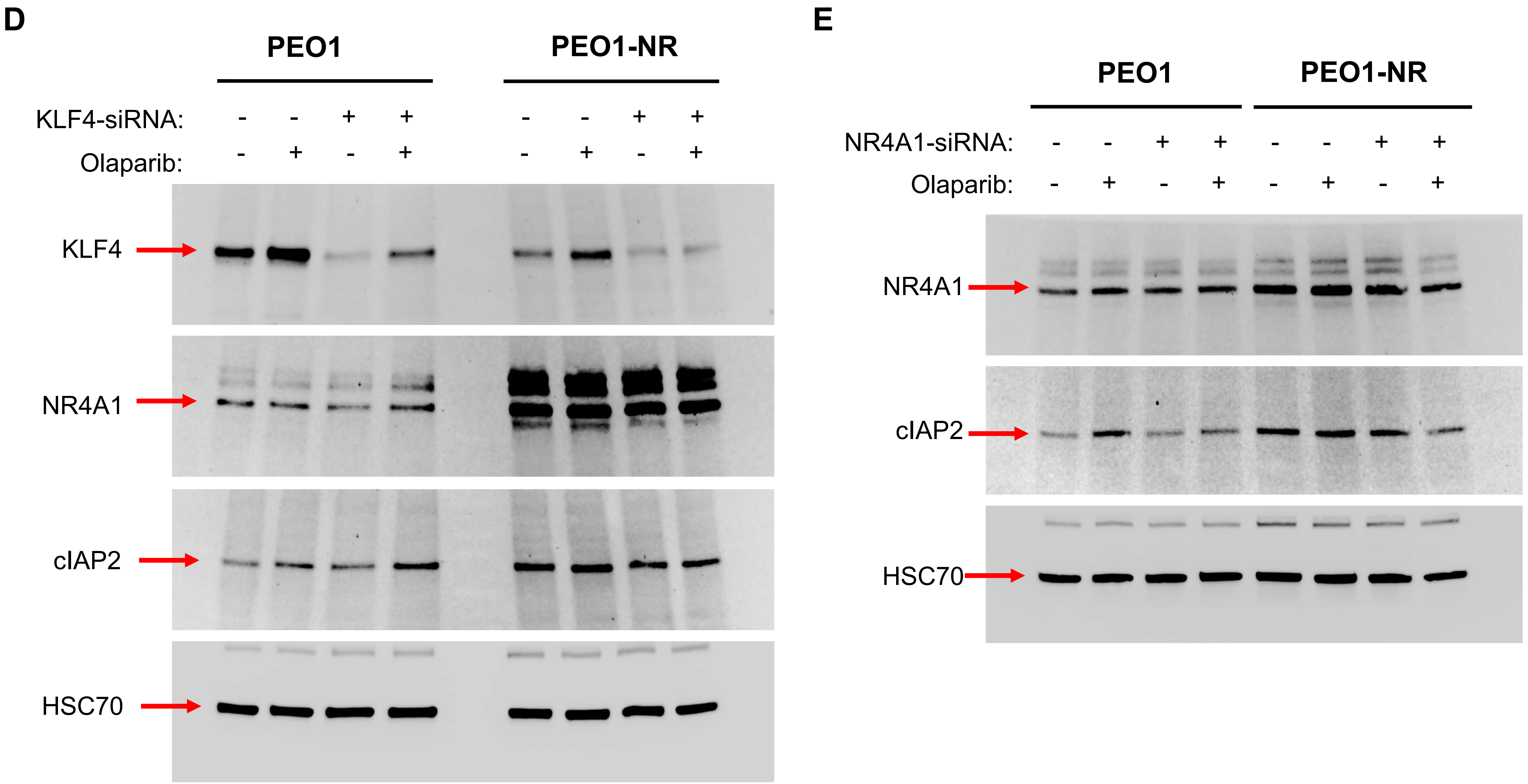


**Supplementary Figure 5-cont.**
